# The bone ecosystem facilitates multiple myeloma relapse and the evolution of heterogeneous proteasome inhibitor resistant disease

**DOI:** 10.1101/2022.11.13.516335

**Authors:** Ryan T. Bishop, Anna K. Miller, Matthew Froid, Niveditha Nerlakanti, Tao Li, Jeremy Frieling, Mostafa Nasr, Karl Nyman, Praneeth R Sudalagunta, Rafael Canevarolo, Ariosto Siqueira Silva, Kenneth H. Shain, Conor C. Lynch, David Basanta

## Abstract

Multiple myeloma (MM) is an osteolytic plasma cell malignancy that, despite being responsive to therapies such as proteasome inhibitors, frequently relapses. Understanding the mechanism and the niches where resistant disease evolves remains of major clinical importance. Cancer cell intrinsic mechanisms and bone ecosystem factors are known contributors to the evolution of resistant MM but the exact contribution of each is difficult to define with current *in vitro* and *in vivo* models. However, mathematical modeling can help address this gap in knowledge. Here, we describe a novel biology-driven hybrid agent-based model that incorporates key cellular species of the bone ecosystem that control normal bone remodeling and, in MM, yields a protective environment under therapy. Critically, the spatiotemporal nature of the model captures two key features: normal bone homeostasis and how MM interacts with the bone ecosystem to induce bone destruction. We next used the model to examine how the bone ecosystem contributes to the evolutionary dynamics of resistant MM under control and proteasome inhibitor treatment. Our data demonstrates that resistant disease cannot develop without MM intrinsic mechanisms. However, protection from the bone microenvironment dramatically increases the likelihood of developing intrinsic resistance and subsequent relapse. The spatial nature of the model also reveals how the bone ecosystem provides a protective niche for drug sensitive MM cells under treatment, consequently leading to the emergence of a heterogenous and drug resistant disease. In conclusion, our data demonstrates a significant role for the bone ecosystem in MM survival and resistance, and suggests that early intervention with bone ecosystem targeting therapies may prevent the emergence of heterogeneous drug resistant MM.

## Introduction

Multiple myeloma (MM) is the second most common hematological malignancy in the US^1^, characterized by the clonal expansion of plasma cells localized primarily in the bone marrow microenvironment of the skeleton ^2,3^. In the bone microenvironment, MM cells disrupt the finely tuned balance of trabecular and cortical bone remodeling to generate osteolytic lesions. This is primarily via MM-induced inhibition of osteoblastic bone formation and increase of osteoclastic bone resorption. There are currently several approved therapies to treat MM including, but not limited to, proteasome inhibitors (e.g., bortezomib and carfilzomib), chemotherapy (e.g., melphalan), immunomodulatory drugs (e.g., lenalidomide), and cellular and non-cellular immunotherapies (e.g., B cell maturation antigen (BCMA) chimeric antigen receptor-(CAR) T cells and daratumumab) that significantly contribute to the long-term survival of MM patients^4^. However, MM remains fatal as nearly all patients ultimately become refractory to various lines of therapy. A greater understanding of how resistant MM emerges in the clinical setting can yield new treatment strategies to slow relapse times and extend the efficacy of standard of care therapies.

In MM and other cancers, the tumor microenvironment is a well-known contributor to the development of treatment resistance^5–10^. MM is an archetype of cancer cell-stroma interactions whereby MM cells reside and metastasize systemically within the bone marrow microenvironment^11^. In the bone ecosystem, MM interacts closely with mesenchymal stem cells (MSCs) and other progenitor cells^6,12–15^ that secrete MM pro-survival and proliferative factors. Additional studies have shown that the bone marrow microenvironment can also protect MM cells from applied therapy in a phenomenon known as environment mediated drug resistance (EMDR)^7,8,16^. Reciprocally, it is well established that MM-derived factors stimulate osteoblast lineage cells to recruit and activate osteoclasts through receptor activator of nuclear factor kappa-Β ligand (RANKL) production, which in turn degrade the bone matrix to release stored cytokines and growth factors (e.g., transforming growth factor-beta, TGF-β) that support MM survival and growth^2,3,5–10^. MM cells also suppress osteoblast maturation, thus tipping the balance towards excessive bone destruction^2,3^. This vicious cycle leads to complications for MM patients such as hypercalcemia and pathological fracture that greatly contribute to morbidity and mortality^17^.

While EMDR can contribute to drug resistance, MM intrinsic mechanisms, such as genetic and epigenetic alterations, also play a significant role in the process. For example, increased expression of the anti-apoptotic BCL2 protein family is important in mediating MM resistance to proteasome inhibition^18–21^. In fact, depending on selective pressures by different lines of therapies and clonal mutation capacity, MM may evolve to develop multiple mechanisms of resistance^5,22,23^. While EMDR and intrinsic drug resistance mechanisms contribute to the emergence of refractory disease several questions remain: i) How does the interplay between each mechanism contribute to the evolution of resistant MM? ii) How does each mechanism contribute to the heterogeneity of the disease under either control or treatment conditions? iii) Could targeting EMDR alter the course of the disease or its responsiveness to applied therapy thereby yielding significantly greater depths of treatment response? In the context of the bone marrow microenvironment and the direct/indirect communications occurring in a temporal and spatial fashion between MM and stromal cells, these questions remain difficult to address using current *in vivo* and *ex vivo* approaches.

Mathematical models of the tumor microenvironment have become a valuable approach that allows us to interrogate the interactions between multiple cellular populations over time under normal or disease conditions. In particular, hybrid cellular automata (HCA) models which couple discrete cell-based models with partial differential equations include spatial aspects enabling users to locate where phenomena are occurring. Further, mathematical models have the potential for clinical translatability and can be used to study the impact of treatment on the tumor microenvironment and delay the onset of resistant disease^24–31^. We therefore leveraged these modeling approaches to examine MM progression and spatial evolution in response to therapy. To this end, we generated a novel HCA model powered by biological parameters from a mouse model of MM and demonstrate how it captures both normal bone turnover, and spatiotemporal cellular dynamics during MM progression in bone, such as loss of osteoblasts, increased stromal cell infiltration, osteoclast formation, and osteolysis. Additionally, we incorporate the proliferative and protective bone marrow microenvironment effects on MM mediated by osteoclastic bone resorption and bone marrow stromal cells. In the treatment setting, the HCA demonstrates that EMDR contributes to minimal residual disease, protecting tumors from complete eradication. Moreover, this reservoir of MM cells increases the likelihood of unique resistance mechanisms arising over time and greatly contributes to the evolution of MM heterogeneity over time compared to non-EMDR scenarios. These results highlight the importance of the bone marrow microenvironment in contributing to MM resistance and patient relapse and provide a strong rationale for targeting both the MM and bone marrow microenvironment mechanisms for optimal treatment response.

## Results

### The HCA model recapitulates the key steps of trabecular bone remodeling

Trabecular bone remodeling is a continuously occurring process that is essential for regulating calcium homeostasis and bone injury repair (Fig. 1a) ^32,33^. This process occurs asynchronously throughout the skeleton at individual sites known as basic multicellular units (BMUs). The BMU in our HCA consists of five different cell types, including precursor osteoclasts, active osteoclasts, mesenchymal stem cells (MSCs), preosteoblasts, and osteoblasts. Additionally, we include central signaling molecules such as RANKL, a driver of preosteoclast motility and OC formation^34^ and osteoclastic release of bone derived factors (BDFs) such as transforming growth factor beta of which bone is a major reservoir^35^. In the HCA, we use TGF-β as a representative BDF given its known biphasic effects on cell behavior and its concentration dependent ability to modulate preosteoblast proliferation and maturation to bone forming osteoblasts^36–40^. We defined the roles and interactions of these key cellular populations and the factors that govern their behavior during the remodeling process using parameters derived experimentally or from the literature (Fig. 1b-c, Supp Fig. 1a and Supplementary Tables 1-6). It is important to note that each cell in the HCA behaves as an autonomous agent that can respond to surrounding environmental cues independently. In our HCA model, new bone remodeling events are initiated over time at randomly selected locations such that multiple BMUs may be present at a given time as opposed to our previous HCA model^27,41–43^. We demonstrated that each BMU undergoes five key phases of bone remodeling: initiation, resorption, reversal, formation, and quiescence^44^ ultimately returning to homeostasis (Fig. 2 and Supplementary Video 1). We also observed that each simulation (n = 25) generated multiple BMUs over time, resulting in the replacement of the majority of the bone over a four-year period (Fig. 2, Supplementary Video 1, Supplementary Fig. 1). We also noted that while the bone area to total area (BA/TA) remains relatively constant the shape of the trabecular bone changes, underscoring the dynamic nature of HCA model and how it reflects the *in vivo* scenario.

**Fig. 1.**
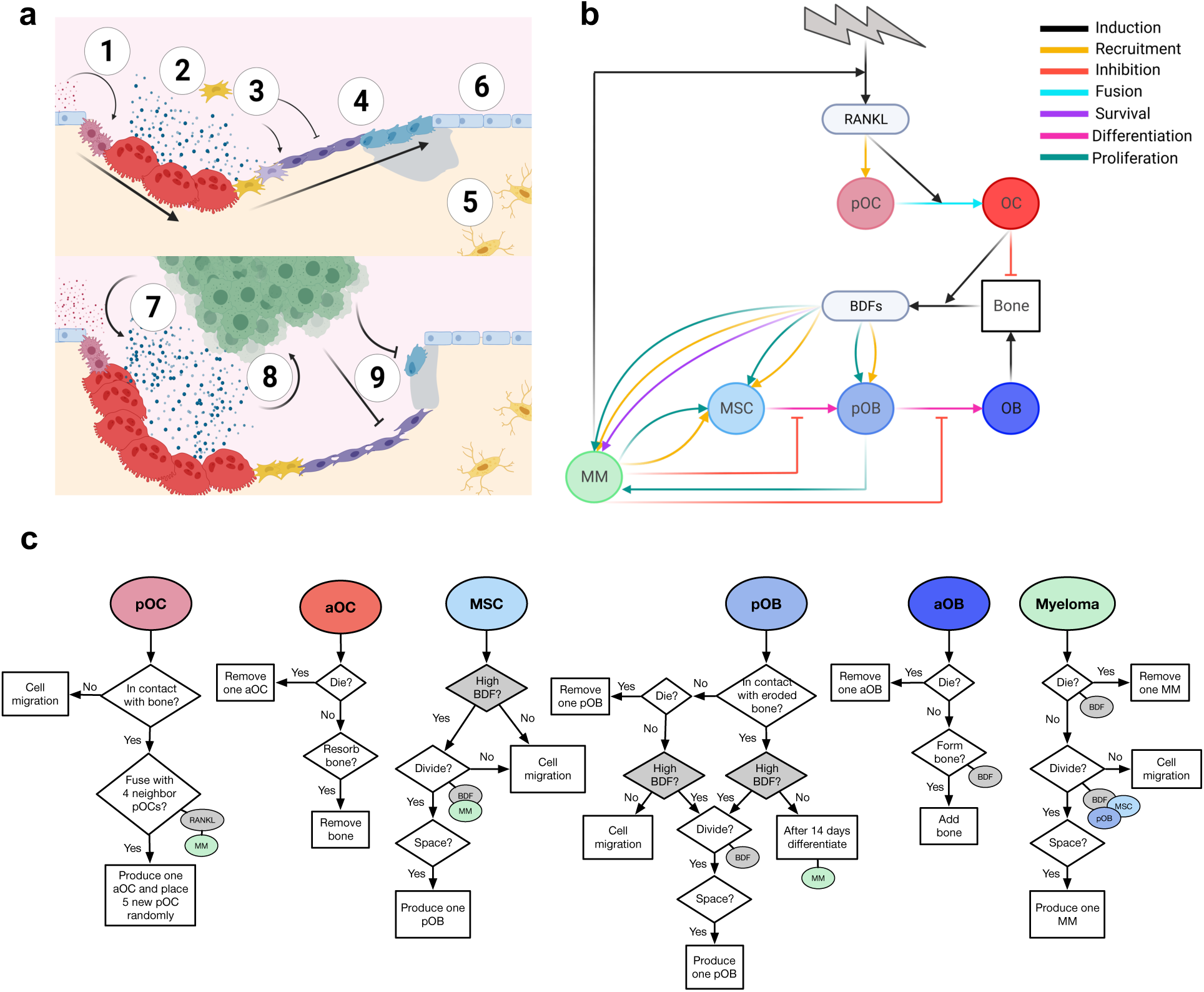
Development of a hybrid HCA of the naïve and myeloma bone microenvironment. **a**, Bone-lining osteoblast lineage cells release RANKL inducing fusion and maturation of osteoclasts (1). Osteoclasts resorb the bone, releasing stored bone-derived factors (BDFs) such as TGF-β (2). TGF-β recruits local MSCs and stimulates asymmetric division in to preosteoblasts (2). When TGF-β levels remain high, preosteoblasts rapidly proliferate. Following osteoclast apoptosis, release of TGF-β falls and preosteoblasts differentiate to mature bone producing osteoblasts (3). While TGF-β levels remain low, osteoblasts produce new bone (4). As bone returns to normal, a fraction of the osteoblasts is buried within the matrix becoming terminally differentiated osteoblasts (5), the remaining osteoblasts undergo apoptosis, or become quiescent bone-lining cells (6). Myeloma cells enhance the formation of osteoclasts (7), enhanced bone resorption produces higher levels of BDFs which fuel myeloma growth (8) and inhibit osteoblast differentiation (8) and activity (9). Created with biorender.com **b**, Interaction diagram between cell types in the HCA and factors such as BDFs and RANKL (created with biorender.com). A more detailed interaction diagram with references can be found in supplementary figure 1. **c**, Flowcharts describing the sequence of steps followed by preosteoclasts, osteoclasts, MSCs, preosteoblasts, osteoblasts, and myeloma cells.

**Fig. 2.**
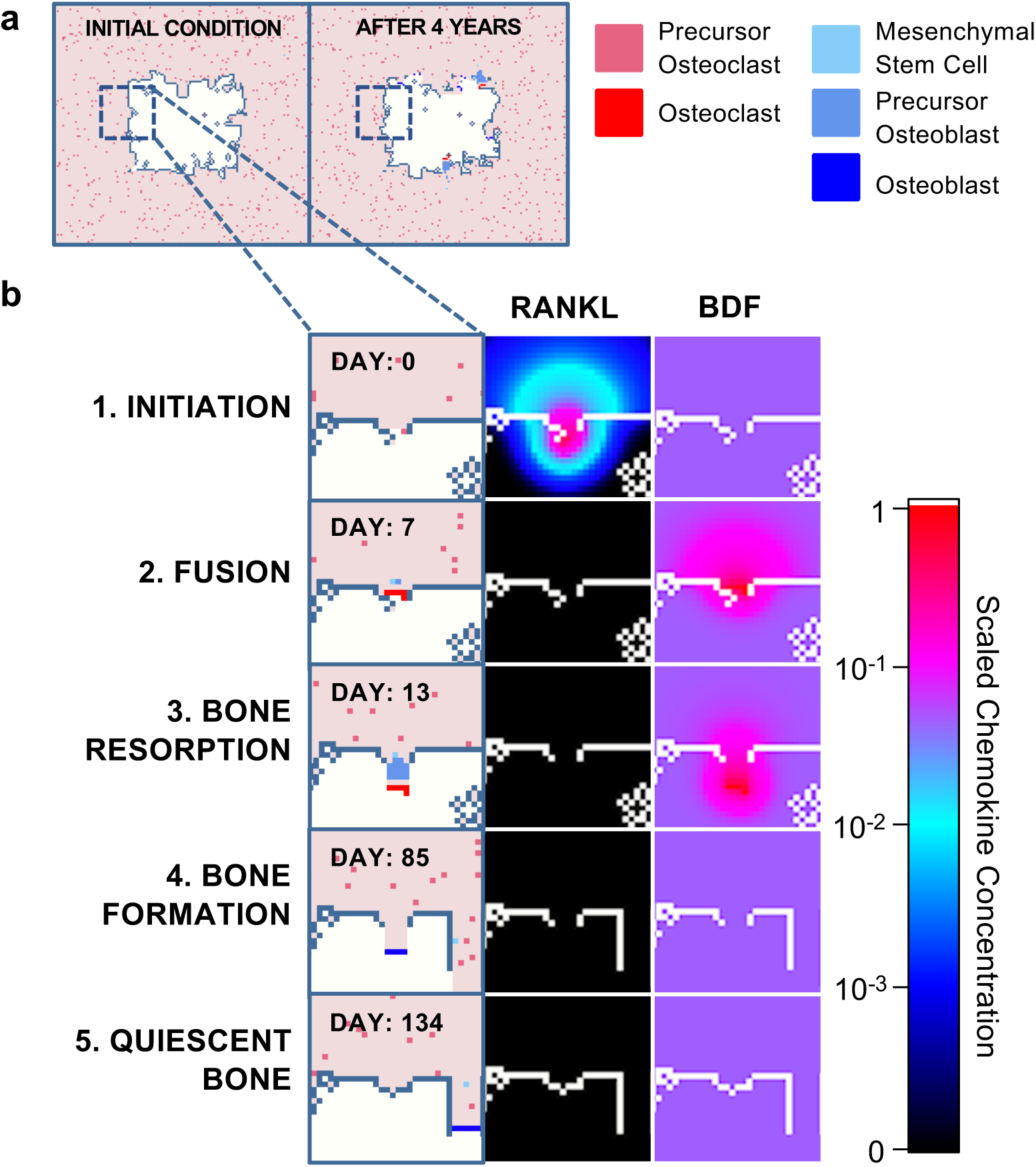
The HCA model captures all stages of bone homeostasis. **a**, Images from simulations at initial conditions (left) and after 4 years (right). Legend depicts colors of cell types in the model. **b**, BMU is initiated in response to release of RANKL from bone lining cells (1). preosteoclasts migrate to RANKL and fuse to form an osteoclast under highest concentrations (2). osteoclast resorbs bone and stored BDFs are released which recruits MSCs. MSCs divide asymmetrically producing preosteoblasts which proliferate rapidly under high TGF-β conditions. Upon completion of resorption, BDF levels fall allowing for preosteoblast differentiation to osteoblasts. osteoblasts form new bone (4) and ultimately undergo apoptosis or become quiescent bone lining cells (5).

### The HCA model captures dose-dependent effects of BDF and RANKL

To challenge the robustness of our bone remodeling HCA, we altered the levels of key factors controlling bone homeostasis. Our *in vitro* experimental data show that neutralizing TGF-β or addition of exogenous TGF-β in osteoblastic cell cultures increases and decreases mineralization respectively, a finding that is consistent with the literature^35,40,45^ (Supplementary Fig. 2a). Similarly, lowering TGF-β levels *in silico* led to a dramatic increase in bone volume (Supplementary Fig. 2b-e). Interestingly, the HCA results showed that when TGF-β levels dropped below 90%, there was significant bone loss (Supplementary Fig. 2b-e) due to abrogation of MSC proliferation leading to reduced numbers of preosteoblasts that contribute to bone formation (Supplementary Fig. 2e). These data are reflective of the known biphasic roles of TGF-β on cell behaviors and processes at high and low concentrations^35,40,45^. In addition, we demonstrated the sensitivity of OC formation in the HCA model in response to changes in RANKL levels (Supplementary Fig. 2f-h). These data demonstrate that the complex HCA model can integrate the behaviors and responses of multiple highly coupled cell types and stimuli to achieve responses reflective of known biology.

### The HCA model integrates the impact of standard of care therapies on bone remodeling

Next, we assessed the response of the HCA model to standard of care therapies used for the treatment of MM such as zoledronate (ZOL) and bortezomib (BTZ). These compounds can block osteoclast function and formation respectively but, as we and others have shown, they can also stimulate osteoblast mediated bone formation^46–49^ (Supplementary Fig. 3a-b and Supplementary Fig. 4a-b). Our results show that, as expected, both ZOL and BTZ therapy increased bone volume over the treatment period by enhancing osteoblastic bone formation and blocking osteoclast activity/formation (Supplementary Fig. 3c-h, Supplementary Fig. 4c-h). While osteoclasts are impacted by both treatments, the HCA recapitulates their distinct mechanisms of action showing that ZOL targets actively resorbing osteoclasts as demonstrated by a reduction in bone resorption per OC (Supplementary Fig. 4h) while BTZ limits the fusion of osteoclast precursors as shown by a reduction in cumulative osteoclast numbers with treatment (Supplementary Fig. 3g). Collectively, these data demonstrate the ability of our HCA model to capture normal bone remodeling over extended periods and the appropriate response of the model to applied therapeutics used for the treatment of MM.

### The HCA model incorporates the MM-bone ecosystem vicious cycle

A key component of MM growth involves the initiation of a feed-forward vicious cycle leading to reduced bone formation and enhanced osteoclastic bone destruction (Fig. 1a). MM is also known to interact with several cellular components of the bone ecosystem such as MSCs^2,16^. To incorporate these effects and MM growth characteristics into the HCA, mice were inoculated with U266-GFP MM cells and at end point, bones were processed and stained for proliferation (Histone H3 phosphorylation (pHH3)) and apoptosis markers (Terminal deoxynucleotidyl transferase dUTP nick end labeling (TUNEL)). Our data show that the highest proportion of proliferating MM cells are located within 50μm of bone (Fig. 3a-b). We noted that fewer cells (TUNEL+) underwent apoptosis in this area (Fig. 3c-d) highlighting the protective effect of BDFs being generated from bone degradation. To consider the effect of bone marrow stromal cells on MM cells distal from bone, we performed *in vitro* experiments and found a significant increase in the proliferation of MM cells grown in the presence of conditioned medium from MSCs and preosteoblasts, but not osteoblasts (Fig. 3e-f). To integrate these observations into the HCA, we assume that the intrinsic rate of MM cell division increases with BDF and when a preosteoblast or MSC is near the MM cells (Fig. 3g-h and Supplementary Equation 11).

**Fig. 3.**
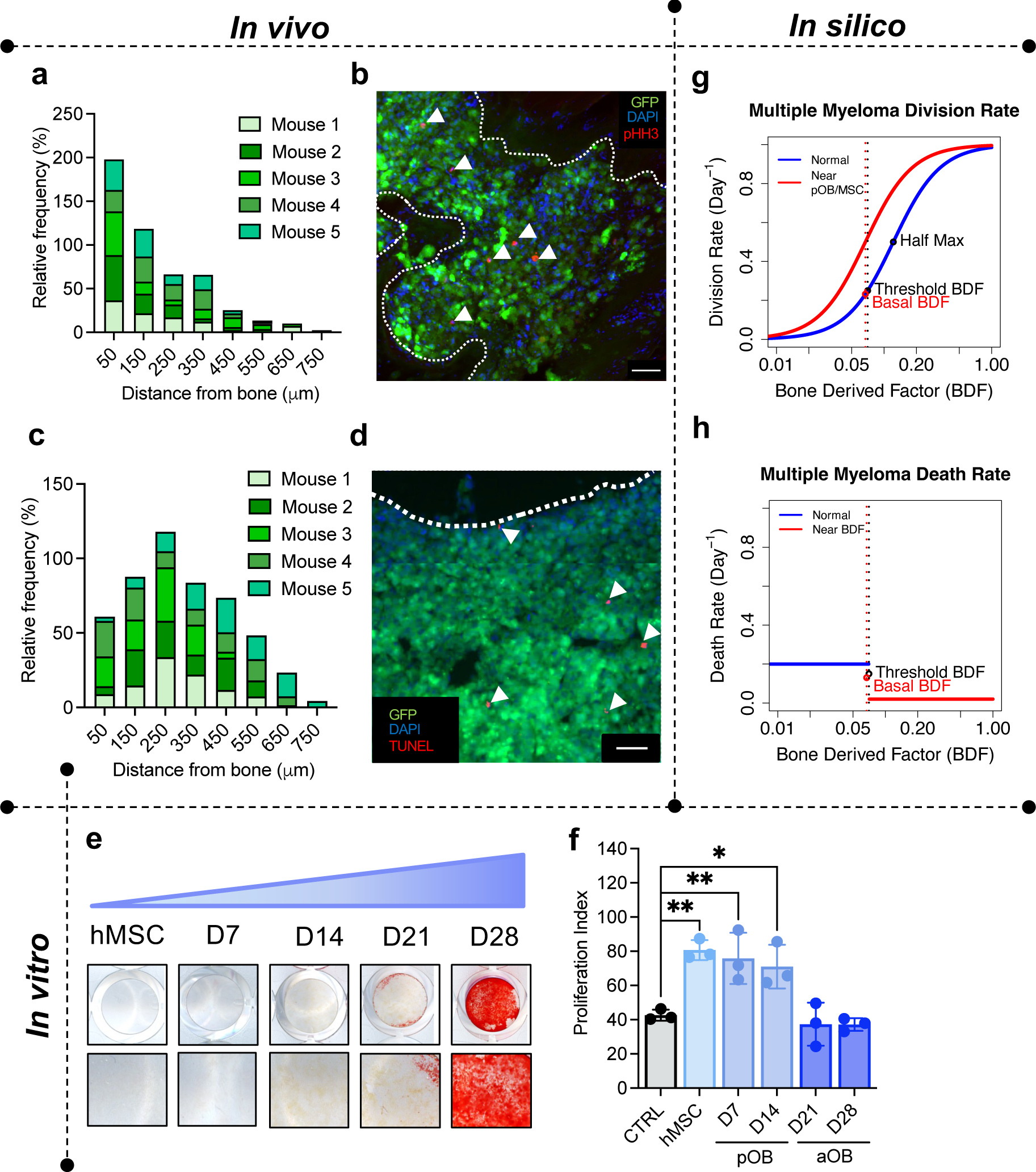
Multiple myeloma cells receive proliferative and survival advantages from the bone marrow microenvironment. **a,** Quantification of the distance of phosphorylated histone H3 positive (pHH3+; red) myeloma cells (green) to the nearest trabecular or cortical bone in U266^GFP^-bearing mice 100 days post inoculation. N=5 tibia. **b,** Representative images from experiment in **a**, DAPI (blue) was used as a nuclear counterstain. White dotted line indicates tumor bone interface. Scale bar, 50 microns. **c,** Quantification of the distance of TUNEL+ (red) myeloma cells (green) to the nearest trabecular or cortical bone in U266^GFP^-bearing mice 100 days post inoculation. N=5 tibia **d,** Representative image from experiment described in **c**, DAPI (blue) was used as a nuclear counterstain. White dotted line indicates tumor bone interface. Scale bar, 50 microns. **e,** Images of huMSCs differentiated to different stages of the osteoblast lineage. Cells were stained with Alizarin Red to identify mineralization. **f,** Proliferation index of CM-DiL stained U266 cells 7 days after growth in 50% conditioned medium from control wells or cells of the osteoblast lineage. Results are displayed as means of at least 3 experiments. **g,** Plot of functional form used to represent the division rate of myeloma cells in the presence and absence of preosteoblasts/MSCs. **h,** Plot of functional form used to represent the death of myeloma cells when BDF is above and below a threshold. * p-value < 0.05, ** p-value <0.01

Next, to study the interactions of the MM-bone marrow microenvironment in the HCA model, we initialized each simulation with a single MM cell near a newly initiated bone remodeling event and the formation of an osteoclast. This assumption is a simplification of the colonization process, in which rare colonizing myeloma cells migrate to the endosteal niche^50,51^. The HCA model revealed exciting temporal dynamics where MM promotes successive phases of bone destruction mediated by osteoclasts that in turn drives MM growth, thereby successfully recapitulating the vicious cycle (Supplementary Fig. 4a and Supplementary Video 2). As expected, the mathematical model demonstrated increasing tumor burden over time (Fig. 4a-b) with rapid loss of bone, correlating inversely with tumor burden (Fig. 4c-d). These findings are consistent with *in vivo* data obtained from the U266-GFP mouse model of MM which showed increasing levels of GFP positive cells in the bone marrow following *ex vivo* flow cytometric analysis (Fig. 4e) combined with a rapid onset of trabecular bone loss (BV/TV; Fig. 4f-g) and associated trabecular bone parameters as assessed by high resolution μCT (Supplementary Fig. 5). The similarities in trends between the *in vivo* model and the HCA outputs, indicated that the HCA model was accurately recapitulating the MM-bone vicious cycle.

**Fig. 4.**
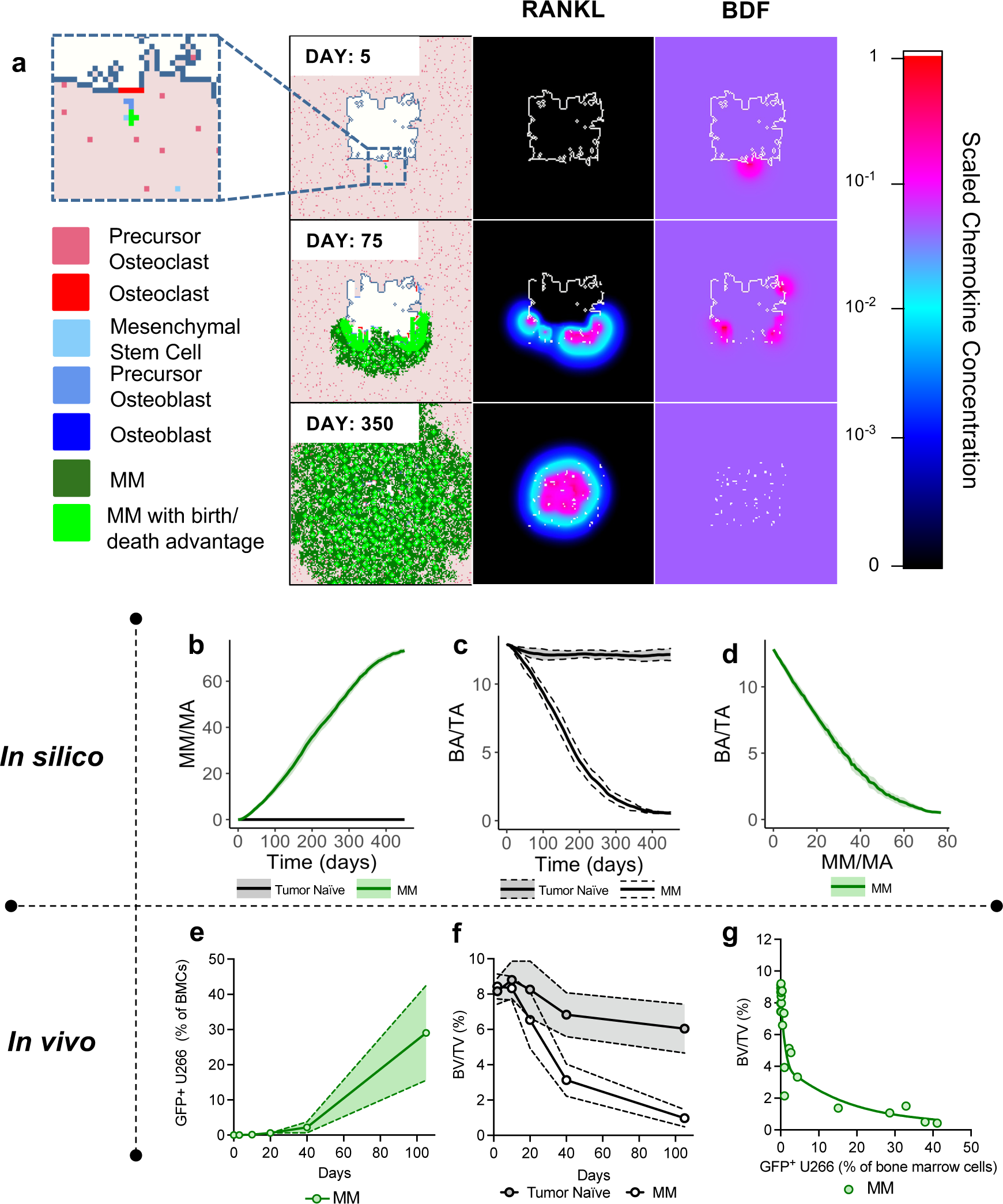
Computational and biological model outputs of myeloma growth and bone dynamics. **a**, HCA model images showing single myeloma cell (Day 5), colonization of the marrow by MM cells (Day 75), increased osteoclastogenesis (Day 75, RANKL), bone resorption (Day 75; BDF) and eventual takeover of the marrow by MM cells (Day 350). Light green myeloma cells indicate MM cells with proliferative or survival advantage. **b**, Myeloma growth dynamics in HCA model in the absence of treatment. **c**, Myeloma induced loss of trabecular bone in HCA model compared to normal bone homeostasis. **d**, Bone loss decreases rapidly with myeloma expansion *in silico*. **e**, Myeloma growth in bone marrow of mice inoculated with U266-GFP cells over time N=3-5 tibia per time point. **f**, Trabecular bone volume fraction (BV/TV) was assessed *ex vivo* with high-resolution microCT. N=3-5 tibia per time point. **g**, Biological model shows rapid myeloma-induced bone loss.

Furthermore, the spatial and temporal nature of the HCA allowed us to make several observations: 1) In the model, the recruitment of MSCs *in silico* increases over time (Fig 5a) as we and others have previously reported^52,53^, 2) the model accurately shows a suppression of adult osteoblast formation (Fig. 5b) due to inhibition of differentiation by MM^2,54^, and 3) the model predicts an early increase in preosteoblasts followed by a reduction in numbers (Fig. 5c), consistent with previous reports^55^, while there is a concomitant increase in osteoclasts (Fig. 5d) ^2,54^. We also compared these population dynamics with our *in vivo* model. In tissue sections derived the U266-GFP model from various time points (days 10, 40 and 100), we measured MSC, preosteoblast, osteoblast, and osteoclast content and observed strong agreement with cell population trends generated by the MM HCA (Fig. 5e-h and Supplementary Fig. 6). Taken together, these data demonstrate the ability of the MM HCA to capture the vicious cycle of MM progression in the bone microenvironment over time.

**Fig. 5.**
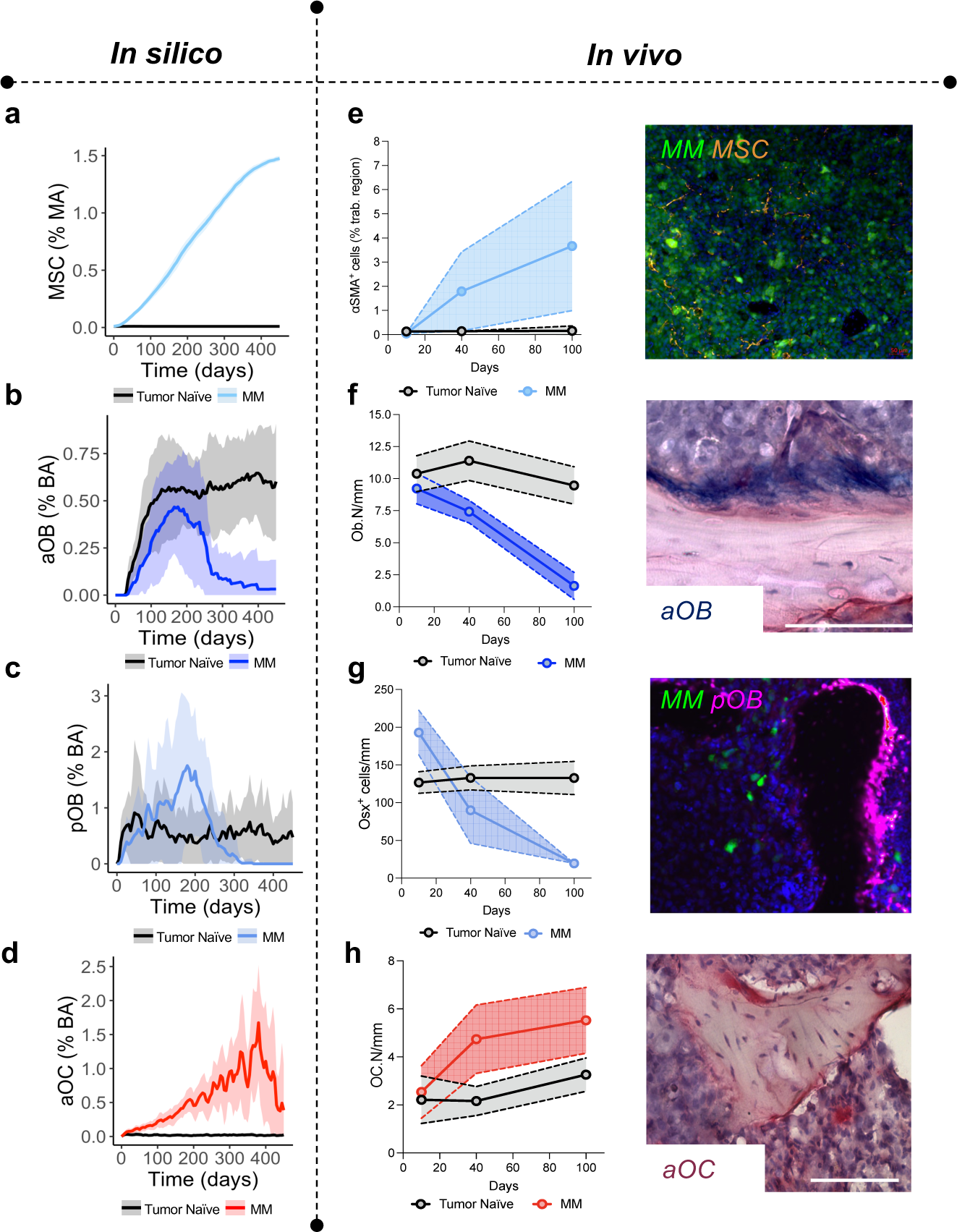
Computational and biological model outputs of cell types in the myeloma bone microenvironment. **a-d,** Computational model outputs of increasing MSC percentage of the marrow area (%MA) (**a**), loss of osteoblasts (**b**) rise and fall of preosteoblast percentage (**c**) and increasing osteoclast percentage due to growth of MM (**d**). **e-h**, *Ex vivo* analysis of histological sections from the U266-GFP myeloma model (N=3-5 tibia per time point.) per time point demonstrates increasing presence of *α*SMA+ MSCs (**e**), loss of ALP+ cuboidal osteoblasts (**f**), early increase and subsequent reduction of OSX+ preosteoblasts (**g**) and increasing numbers of TRAcP+ multinucleated osteoclasts (**h**) compared to tumor naïve mice.

### Environment-mediated drug resistance increases minimal residual disease and leads to higher relapse rates

We and others have shown that BTZ demonstrates a dose dependent cytotoxic effect on MM (Supplementary Fig. 3a)^56,57^. The compound is also effective *in vivo* but MM still grows albeit at a significantly slower rate^56^. Patients also typically become refractory to BTZ. This indicates that MM cells are 1) protected by the surrounding bone ecosystem, or 2) due to selective pressure and mutation or other mechanisms such as epigenetic changes, drug resistant clones emerge, or 3) that both ecological and evolutionary contributions can occur in parallel. Here we sought to use the spatial nature of the HCA model to gain a deeper understanding of how MM resistance evolves focusing initially on the contribution of bone ecosystem, i.e., EMDR. We confirmed the EMDR protective effect conferred by bone stromal cells on MM in the presence of BTZ *in vitro* (Fig. 6a) and integrated this effect into the HCA model (Fig. 6b). BDF was also included since it can provide a survival advantage to MM cells in the presence or absence of treatment^58,59^. Next, we initialized HCA simulations with a homogeneous sensitive MM population. BTZ was applied once the tumor burden reached a volume of 10% to mimic the scenario of diagnosis in the clinical setting. Our results show that without EMDR, BTZ completely eradicated the disease at high doses (Fig. 6d, Supplementary Fig. 3j), consistent with our *in vitro* findings. However, when EMDR effects are included in the model, BTZ reduced MM burden significantly, but the disease persisted at a stable, albeit lower, level and did not grow over time compared to existing *in vivo* data (Fig. 6c-d). These data suggest that EMDR does not directly contribute to MM relapse but does protect a small volume of MM cells from BTZ. Therefore, to address the relapse and the emergence of BTZ resistance in MM cells, we integrated a resistance probability (pΩ = 10^−4^ or pΩ= 10^−3^) during cell division, which allowed for the emergence of BTZ resistance. The probability of a MM cell to develop resistance to BTZ are estimated based on the current literature, in which mammalian cancer cells have a probability of developing resistance to numerous treatments of between 10^−3^ to 10^−6 60,61^. Based on our *in vitro* observations, we also incorporated a cost of resistance^62^ in the model such that BTZ-resistant cells proliferate at a slower rate and are out-competed by PI-sensitive cells in the absence of treatment (Supplementary Fig. 7).

**Fig. 6.**
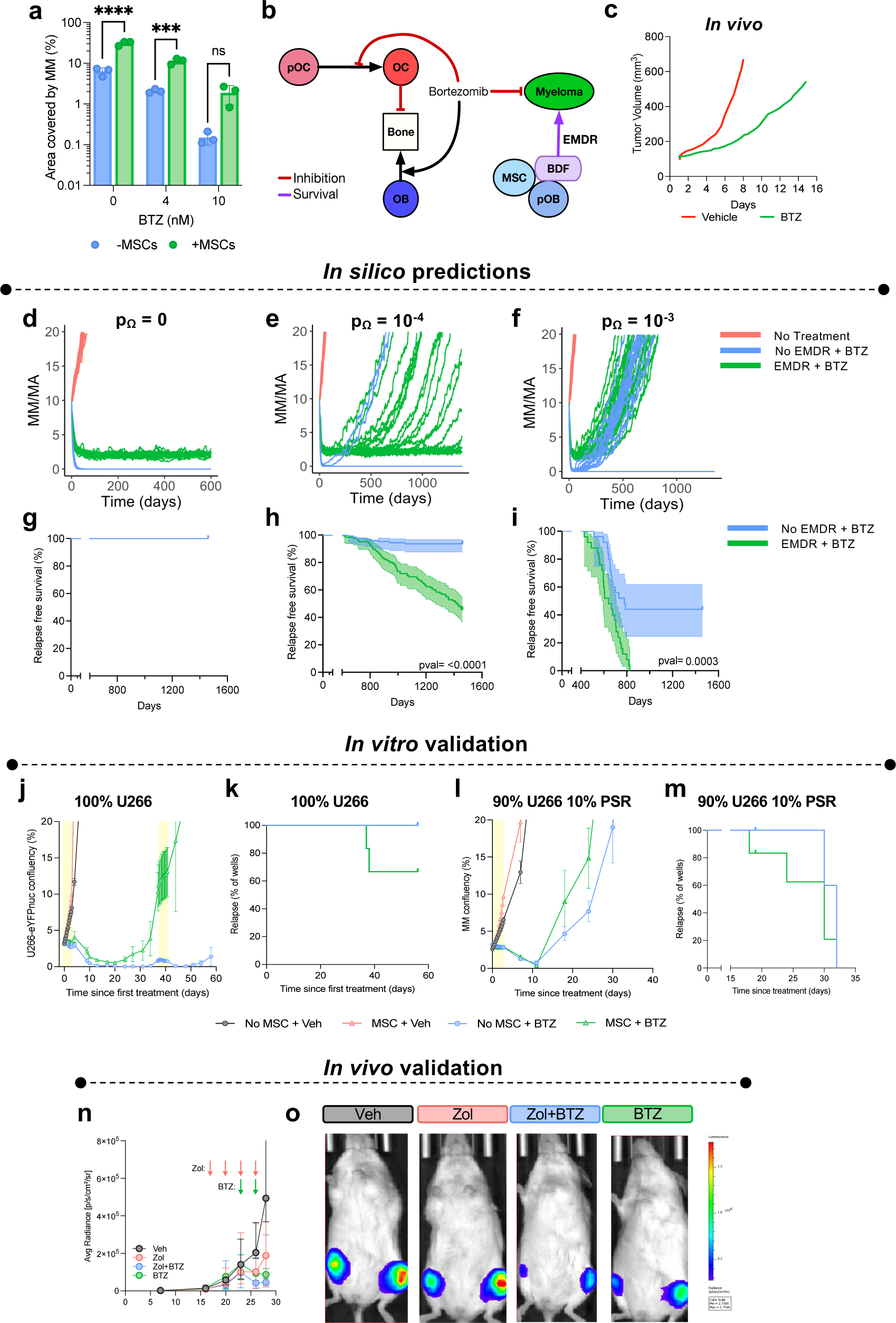
EMDR contributes to minimal residual disease and relapse. **a,** GFP+ U266 MM were plated alone or with huMSCs in the presence or absence of BTZ; MM burden was measured after 72h by area covered by GFP+ cells. **b**, Interaction diagram showing the cell types and factors in the HCA that are affected by BTZ or contribute to EMDR. **c**, *In vivo* growth of MM cells with and without BTZ treatment, data was taken from ^42^. **d-f,** Model outputs of MM growth with continuous BTZ treatment, +/− EMDR when MM cells do not develop resistance (**d**) or with a probability to develop resistance (10^−4^, **e** or 10^−3^, **f**) and develop resistance. **g-i**, Kaplan-Meier plot of relapse-free survival from simulations described in **d-f**. **j,** Nuclear-eYFP+ U266 were cultured alone or with huMSCs in the presence or absence of 10nM of BTZ. MM growth was tracked by eYFP over 60 days. Yellow shading indicates periods when treatment was on. **k**, Kaplan-Meier plot of ‘relapse’ (wells reaching >20% MM confluency) from experiment described in j. **l**, MM cells (90% U266-Nuclear-eYFP+ and 10% PSR-RFP+) were cultured alone or with huMSCs in the presence or absence of 10nM of BTZ. **m**, Kaplan-Meier plot of ‘relapse’ (wells reaching >20% MM confluency) from experiment described in l. **n**, Average U266 growth by BLI after MM cells (90% U266-GFP^+^Luc^+,^ 10% PSR-RFP) were tail vein injected into NSG mice. Mice were divided into two groups and pre-treated with vehicle or Zol (30ug/kg) for 1 week prior to mice being randomized and treated with either vehicle (n=5 mice), Zol (n=5 mice), Zol+BTZ (n=7 mice), or BTZ (0.5mg/kg; n= 4 mice). Pink and green arrows indicate days of Zol or BTZ treatment respectively. **o**, Representative BLI IVIS images from the experiment described in **n**.

We then repeated our *in-silico* experiments and observed that with the lower resistance probability (p_Ω_ = 10^−4^), EMDR increased the likelihood of tumor progression with more simulations likely to relapse (54.4%, n = 125) compared to tumors without EMDR (6.4%, n = 125; Fig. 6e and g; Supplementary Video 3). Similarly, with a higher resistance probability (p_Ω_ = 10^−3^) tumors were more likely to relapse when EMDR is present (100%, n = 25) compared to tumors without EMDR (56%, n = 25, Fig. 6f and h; Supplementary Video 4). However, the higher resistance probability also increased the likelihood of tumor progression regardless of EMDR, implying a greater role for MM intrinsic resistance mechanisms during tumor relapse. To strengthen these findings, we tested a more clinically relevant treatment strategy, in which BTZ therapy was pulsed (2 weeks on, 1 week off) and observed similar results (Supplementary Fig. 8). To validate our *in silico* results, we utilized both *in vitro* and *in vivo* methodology. To this end, U266 MM cells were cultured alone (monocultures) or with protective human MSCs (huMSCs) and exposed to vehicle or BTZ. BTZ-treated U266 groups, responded to BTZ as evidence by a reduction in MM confluency (Fig. 6j). However, we observed that BTZ was less efficient when U266 were cultured with huMSC and recovered over a 30-day period, whereas monocultures did not. Further, co-cultured U266 cells treated with a second dose of BTZ, continued to proliferate, even beyond initial confluency, and thus were considered “relapsed”, whereas this was not the case in monocultures (Fig. 6K). Since the propensity of MM cells to develop resistance cannot be readily manipulated, we repeated the *in vitro* experiments with a mixed population of MM cells that were either sensitive (U266; YFP positive) or resistant (U266-PSR; RFP positive). In cultures composed of 90% PI-sensitive U266, 10% PI-resistant U266-PSR, we observed that MSCs protect against BTZ, leading to enhanced relapse compared to MM cells cultured alone thus confirming our *<u>in silico</u>* observations. Next, we investigated the role of EMDR *in vivo*. Currently, there are no approved stromal targeting therapies available for the treatment of MM, however, Zol is frequently given to patients to inhibit osteoclast-mediated bone destruction that in turn also reduces the release of BDF that promote MM survival. NSG mice were inoculated with 90% PI-sensitive U266-GFP-Luc and 10% PI-resistant-RFP MM cells. Upon detection of the tumors in the hindlimbs, mice were divided in to two groups and pretreated with either vehicle or Zol for one week to limit osteoclast-mediated release of BDFs. Following pretreatment, mice were further subdivided and treated with BTZ only or BTZ and Zol for another week. As predicted by our *in-silico* results, limiting EMDR with Zol led to a deeper response to BTZ in PI-sensitive MM, as evidence by bioluminescence imaging and end point flow cytometry (Fig 6n-o).

Our HCA model revealed during BTZ application, EMDR protected a reservoir of sensitive cells that ultimately develops intrinsic resistance leading to the increased relapse rates observed (Fig. 8a-b; Supplementary Fig. 9a). The role of this sensitive reservoir in relapse is particularly evident at the lower resistance probability, where tumors with EMDR exhibited more variable (generally longer) relapse times compared to tumors without EMDR. When EMDR is absent, many tumors go extinct and relapse only occurs if intrinsic resistance arises early following treatment, whereas with EMDR, sensitive cells persist, and resistance can arise later in the course of the disease. We validated this finding *in vitro,* where we demonstrate that MSCs protect a small portion of PI-sensitive U266 MM cells from BTZ treatment, whereas in cultures containing only MM cells, PSR-RFP MM cells ultimately became the sole clone upon relapse following BTZ treatment (Fig 7c-d). Further, *in vivo*, treatment of mice with BTZ, led to the outgrowth of PI-resistant PSR-RFP. However, in Zol-pretreated (EMDR-inhibited) mice we see a deeper reduction in PI-sensitive MM cells compared to BTZ-only treated mice, where EMDR is intact (Fig. 7e-f). We next determined the clinical relevance of whether PI sensitive MM cells existed in patients that were considered to have relapsed refractory MM (RRMM). We isolated CD138+ cells from MM and exposed the cells to BTZ *ex vivo*. We observed that approximately 20% of relapsed refractory MM patients who a failed a PI-containing regimen (n=86) contained MM cells that are still sensitive to PI treatment thus supporting our *in silico* and *in vitro* and *in vivo* findings. Taken together, these results demonstrate the ability of the HCA to predict the effect of resistance and EMDR on tumor relapse and indicate a vital role of EMDR in contributing to minimal residual disease and the development of relapse refractory MM.

**Fig. 7.**
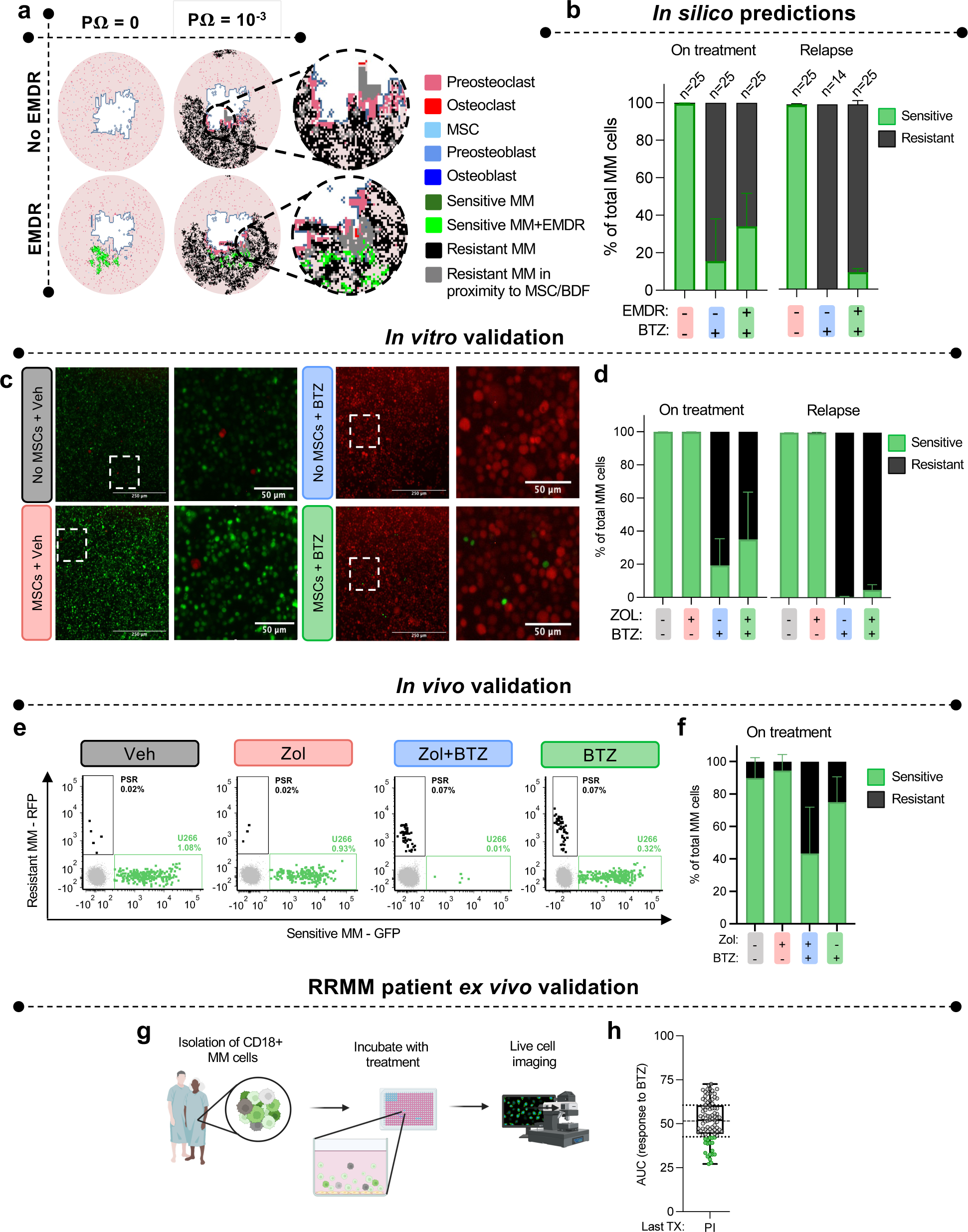
EMDR contributes protection of sensitive MM and tumor heterogeneity. **a,** HCA images from simulations with continuous BTZ application, with no resistance probability (p_Ω_=0) or high resistance probability (p_Ω_=10^−3^) in the presence or absence of EMDR. **b**, Computational outputs of the proportion of BTZ-sensitive and BTZ-resistant MM cells 1 year on treatment (left) and at relapse (right) with p_Ω_=10^−3^. Related to Figures 6f and 6i. **c**, Representative images of U266-nuclear eYFP+ and PSR-RFP+ cells at relapse when cultured alone or huMSCs, with or without BTZ in vitro. Related to figure 6j-k. **d**, The proportion of BTZ-sensitive (U266-nuclear eYFP+) and BTZ-resistant (PSR-RFP+ MM cells cultured alone or with huMSCs with and without BTZ on treatment (left) and at relapse (right). Related to Figures 6l-m and 7d. **e**, Representative flow plots of BTZ-sensitive (U266 GFP+Luc+; green) and BTZ-resistant (PSR-RFP+; black) from the bone marrow (gray) of NSG mice treated with vehicle, Zol, Zol+BTZ or BTZ. Vehicle = 10 femur, Zol = 10 femur, Zol+BTZ = 12 femur, BTZ = 8 femur. **F**, Proportion of BTZ-sensitive (U266 GFP+Luc+) and BTZ-resistant (PSR-RFP+) MM cells in the bone marrow of NSG mice treated with vehicle, Zol, Zol+BTZ or BTZ. **g,** schematic of the EMMA platform. CD138+ MM cells and stroma are isolated from MM patients and co-cultured without test compounds. Live cell imaging is used to assess viability. **h**, The AUC of MM cells from RRMM patients, whose last relapse was to a PI-containing regimen (PI) or a non-PI containing regimen (Other), in response to BTZ. Green dots identify sensitive MM cells (quartile 1).

**Fig. 8.**
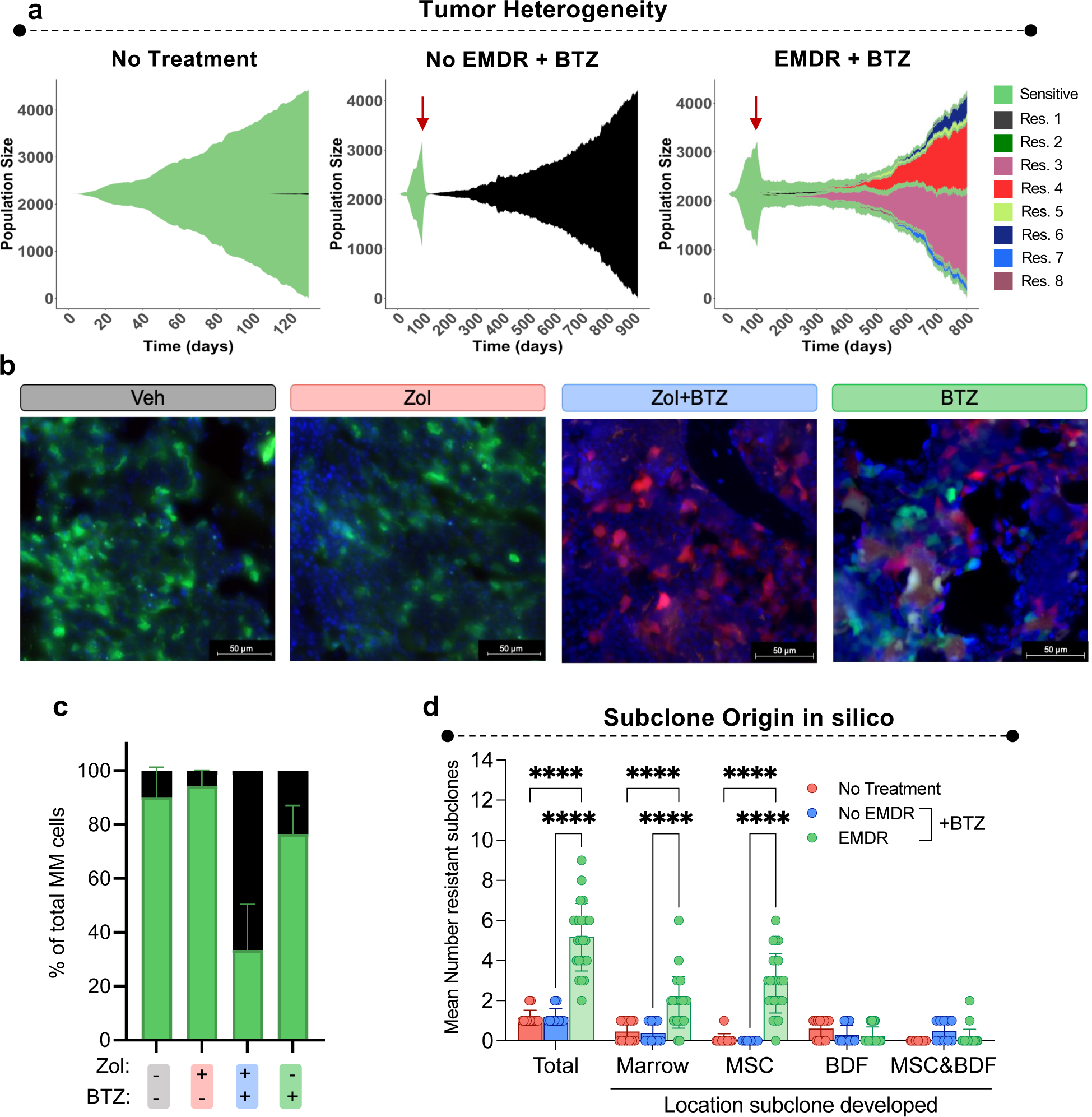
EMDR contributes to tumor heterogeneity upon relapse. **a**, Muller plots of sensitive and individual BTZ resistant (Res) sub-clones from simulations described in **a-c**. Colors denote different subclones. Red arrow indicates start of BTZ treatment. **** pvalue <0.0001 **b**, Images of GFP+ U266 (green) and RFP+ PSR (red) MM cells and cell nuclei (DAPI; blue) in tibial sections of mice from *in vivo* study described in Figure 7. Magnification 20X. Scale bar 50 microns. **c**, Relative quantification of GFP+ U266 and PSR MM in tibial sections described in Figure 8b. Vehicle = 5 tibia, Zol = 5 tibia, Zol+BTZ = 7 tibia, BTZ = 4 tibia. **d**, Mean number of resistant sub-clones arising following BTZ treatment tumors that reached 20% with resistant subclones and the locations within the bone marrow microenvironment where each subclone originated, with/without EMDR from simulations described in **a**. n = 13/25 (No Treatment), n = 10/25 (No EMDR+BTZ), n = 24/25 (EMDR+BTZ).

### Environment-mediated drug resistance and resistance probability impact MM heterogeneity in relapsed disease

Even though relapsed tumors with a high resistance probability had similar growth dynamics with and without EMDR, we noted that the composition of the tumors differed greatly. Myeloma is a clonal disease with the prevalence of clones changing throughout treatment^23,50,63,64^. Given the spatial resolution of the HCA model, we next asked whether EMDR impacted the heterogeneity of resistant tumors. To this end, we tracked the number and spatial localization of new clones forming from unique resistance events under BTZ treatment conditions in the presence or absence of EMDR. We observed that at a high resistance probability (p_Ω_ = 10^−3^) EMDR drove the evolution of significantly more resistant subclones (mean = 5.2 subclones with >10 cells, n = 24 simulations) compared to non-EMDR conditions where the arising population was composed largely of a single resistant clone (mean = 1.2 subclones with >10 cells, n = 10 simulations; Fig. 8a), with similar results at lower resistance probabilities and pulsed BTZ treatment (Supplementary Fig. 9b-c). *In vivo*, we observed a greater reduction in the ratio of GFP+ sensitive to RFP+ resistant MM when mice were treated with Zol+BTZ compared to BTZ only (Fig 8b-c). Moreover, the HCA revealed a significant proportion of resistant subclones originated in close proximity to MSCs/preosteoblasts (Fig. 8d) or following release of BDF from osteoclastic bone resorption (Supplementary Fig. 9c). Taken together, these data would indicate that EMDR not only leads to enhanced relapse, but increased MM heterogeneity due to the emergence of independent resistant MM clones. Further, it is also plausible that EMDR may give rise to MM clones with potentially unique intrinsic BTZ resistance mechanisms making the treatment of relapsed tumors more challenging. Together, these data support that early targeting of bone marrow microenvironment, and thus EMDR, would lead to reduced MM heterogeneity and potentially deeper responses with subsequent lines of treatments.

## Discussion

Understanding the factors that govern the evolution of refractory MM remains of huge clinical importance. Dissecting how intrinsic and bone ecosystem mechanisms contribute to the evolution of the disease in the presence or absence of treatment or where these clones arise spatially are difficult questions to address with standard experimental techniques alone. Here, we address this using a novel experimentally derived hybrid cellular automaton (HCA) model of MM evolution in the bone ecosystem that provides new insights. For example, we show that the bone ecosystem contributes to minimal residual disease, and MM clonal heterogeneity. These findings are in part supported by previous studies revealing enrichment of binding proteins and chemokine receptors on MM cells in minimal residual disease, indicating a likely role for stromal interactions in facilitating relapse^65^. Our findings and additional wider biological validation provide rationale for the targeting of the bone ecosystem during the application of standard of care treatment to prevent the emergence of multiple resistant clones.

Our own *in vitro* and *in vivo* data, as well as data from the literature, were used where possible to parameterize and calibrate the HCA model. However, there were some parameters that could not be estimated, and the specific assumptions that have been incorporated into the HCA may impact the generality of the results. For example, we assume that MSCs are recruited in proportion to tumor burden based on experimental data in non-treatment conditions, but the application of treatment could alter this recruitment rate and thus bone ecosystem protective effects. Furthermore, there are limitations to how well parameters estimated from *in vitro* data correspond to the *in vivo* setting. To address this, we tested the robustness of the model results by varying a subset of parameter values to determine the impact on simulation results (Supplementary Methods 1.4; Supplementary Fig. 10). Overall, our finding that EMDR increases minimal residual disease and leads to higher relapse rates was consistent across parameter values, underscoring the robustness of the model.

There have been several mathematical models that explore interactions between MM and the bone marrow microenvironment. For example, ordinary differential equation (ODE), evolutionary game theory (EGT), and hybrid agent-based models have revealed insights into MM growth in bone and have even taken into consideration the impact of MM progression on bone destruction^66–72^. Additionally, mathematical models have focused on how ecosystem effects contribute to drug resistance^73–76^. However, the HCA model presented herein represents a significant advance in that it is carefully integrated with biological data to recapitulate the bone-MM ecosystem under normal and treatment conditions and is the first model to dissect intrinsic vs. ecosystem driven resistance in multiple myeloma. A major additional advantage of our model is its spatial nature, allowing us to visualize and measure where the ecosystem, and particularly stromal cells, contribute to the generation of resistant clones.

Our model shows that resistant MM clones can arise independently of the bone ecosystem during BTZ treatment if the resistance probability is sufficiently high. But, when EMDR is present, we observe the evolution of higher numbers of resistant clones near MSCs/preosteoblasts/BDF during continuous BTZ treatment. The model also suggests that targeting of EMDR during treatment would reduce MM heterogeneity. Various adhesion molecules and cytokines such as the CXCL12/CXCR4 and IL-6/JAK/STAT axis have been shown to mediate ecosystem protective effects and drugs/biologics that specifically target those pathways have been trialed^77,78^. Unfortunately, these drugs have had little success translating to clinical practice due to limited survival improvements when added to existing therapies^79^, thus there remains a need to identify multiple mechanisms of EMDR in patients with novel therapies and feasible methods of delivering these drugs. However, we posit that early intervention with EMDR-targeting agents, perhaps in combination, would reduce tumor heterogeneity, rendering relapsing tumors vulnerable to a targeted second line therapy that targets the predominantly homogenous disease. As such, future studies/trials should take into consideration the tumor diversity upon relapse and response to second line therapies.

Another important aspect of the model is the ability to apply and withdraw therapy at varying stages of MM progression to quantitatively determine the impact on MM over time which may be useful for adaptive therapy design and implementation. Adaptive therapy is an emerging, evolution-inspired approach for cancer treatment that exploits competitive interactions between drug sensitive and drug resistant cells to maintain a stable tumor burden^62^. Several models have studied the many ways therapies can be applied in an adaptive manner ensuring that the competition between treatment resistant and naïve populations allows for better overall survival for cancer patients. Importantly, our model captures tumor interactions with the bone ecosystem and thus includes the potential impact of the bone ecosystem on adaptive therapy. This allows us to study not only how treatments impact tumor populations but also their influence on bone ecosystem cells such as osteoblasts and osteoclasts, and how this can then influence MM response since the protective bone microenvironment can serve as a reservoir of sensitive cells under treatment. Clinical trials thus far using adaptive therapy guided by mathematical modeling have had encouraging results. For instance, recent studies have shown that in patients with bone metastatic prostate cancer, adaptive therapies guided by mathematical models can increase the time to progression and the overall survival using standard of care treatments^80^. These data support the feasibility of integrating mathematical modeling for the design of patient specific treatment. It should be noted however, the HCA model is parameterized with a variety of in vitro, in vivo, and human data, making translation directly to the clinic challenging in its current form. This can be resolved with further rigorous model calibration and validation using clinical data, and multiple efforts at moving agent-based models into the clinical setting are underway^29–31,81,82^.

A key component of our HCA model lies in its ability to recapitulate the homeostatic nature of bone remodeling over prolonged periods (∼4 years). As we demonstrated, the impact of therapies on tumor-naïve bone volume and the bone stromal cellular components that control the process can also be quantitatively examined. It also can examine non-cancerous bone diseases that result from an imbalance in bone remodeling such as osteoporosis^83^. The underlying causes of osteoporosis can result in differing effects on osteoclasts and osteoblasts, which could be incorporated into the model. For example, estrogen deficiency increases the lifespan of osteoclasts while having the opposite effect on osteoblasts, whereas age-related senescence is characterized by decreased osteoclastogenesis and osteoblastogenesis^83^. Osteoporosis treatments include anti-resorptive therapies (e.g., bisphosphonates) and bone formation therapies (e.g., intermittent PTH). By modifying certain model parameters, we can explore the effect of different therapeutic strategies on disease outcomes to determine if certain treatments are more effective in particular scenarios^84^. While we have primarily used Zol and BTZ for proof of principle, the model can be easily adapted to integrate the effects of other standard of care therapies as single agents or in combination such as dexamethasone, and IMIDs. Importantly, the temporal and cellular resolution of the model also can identify key moments in which to administer treatments to maintain a healthy bone volume.

Our HCA model, like all models, has caveats. For example, its two-dimensional nature does not take into account the potential three-dimensional role in evolutionary dynamics^85,86^ or the vascular nature of the bone marrow-MM microenvironment that can play a role in drug/nutrient diffusion,^10,87^. However, these can be incorporated into the model, if necessary, upon availability of relevant experimental data. Another potential and future extension of the model is to incorporate the immune ecosystem given its role in controlling bone turnover and cancer progression^10,87^, the influence of BDFs on immune populations^88,89^ and the growing armamentarium of immunotherapies in MM. Here, we excluded the immune component to focus solely on the role of the bone stroma on MM progression combined with the fact that MM growth in immunocompromised mice was used to parameterize and compare model outputs. Infiltrating immune cells have noted effects on cancer progression. For example, initially, natural killer (NK) cells and cytotoxic T lymphocytes can drive an anti-tumor response^90^. However, as the tumor progresses, immunosuppressive populations expand including myeloid derived suppressor cells (MDSCs) and regulatory T-cells (Tregs)^90^. MM is characterized by an increase of inactive NK cells, MDSCs, and Tregs and treatments such as bortezomib can alter the composition and activity of infiltrating or resident immune cells^90,91^. Tumor-immune dynamics in MM have been previously explored with ODE models^92,93^ and parameters derived from *in vitro* or *in vivo* studies can be integrated into our HCA model albeit with a limit on complexity. Another caveat is the difficulty in determining the rate of MM resistance and the assumption that it occurs intrinsically due to treatment pressure. Prior studies suggest that drug resistance may arise through the selective expansion of pre-existent resistant tumor cells, *de novo*, or through the gradual increase in resistance over time^94^. In our model, we assume that after treatment initiates (i.e., once the bone marrow consists of 10% of MM cells), there is a probability that a dividing MM cell could develop a resistance mechanism, and that mechanisms directly causes resistance to bortezomib resulting in heterogeneity. This resistance probability that would lead to treatment resistance is difficult to assess in patients or in experimental models but the resultant heterogeneity arising from different resistance probabilities may allow us to infer resistance rates in patients. Moreover, the *in silico* model is not fixated on specific resistance mechanisms currently but more on whether a cell is sensitive or not. One such benefit of the *in silico* model is the ability to alter parameters, such as p_Ω_, response to treatment and doubling time, to mimic the heterogeneity seen in the clinic.

In conclusion, we have described a new mathematical model of MM evolution in the context of the bone ecosystem. Our results show that, under treatment conditions, the interactions between the bone ecosystem and the tumor contributes to the presence of residual disease and to the enrichment of tumor heterogeneity. These results suggest that early intervention with drugs that target the bone ecosystem could lead to MM extinction or at the very least reduce tumor heterogeneity by preventing the evolution of multiple drug resistant clones. Given the current interest in novel strategies of treatment that consider the evolutionary dynamics in cancer such as adaptative therapies, we also predict that our model can be a useful clinical tool in guiding patient specific treatment strategies.

## Methods

### Hybrid Cellular Automaton Model

The model we developed builds on the HCA paradigm first described by Anderson *et al.* and used to study evolutionary dynamics in cancer and our work in the context of bone metastasis^41–43,95,96^. By definition, an HCA consists of discrete cell types that are updated once per time step according to a set of flow charts (Fig. 1c) as well as a set of partial differential equations that describe cytokines in the microenvironment. Here, we consider six different cell types, including precursor osteoclasts, active osteoclasts, mesenchymal stem cells (MSCs), precursor osteoblasts, active osteoblasts, and multiple myeloma cells. Additionally, we incorporate two signaling molecules including RANKL and bone derived factors (BDFs) such as transforming growth factor beta (TGF-β). In the sections below, we describe the interactions between cell types and cytokines that are key in bone remodeling and MM progression. Further details can be found in Supplementary Methods 1.

#### Parameters and grid

We implemented our model using the Hybrid Automata Library ([HAL](https://journals.plos.org/ploscompbiol/article?id=10.1371/journal.pcbi.1007635))^97^. When possible, parameters for the HCA model were derived from empirical and published data (Supplementary Tables 1-6). The model is defined on a 2D rectangular grid (160 × 150 pixels) representing a 1600 × 1500 µm^2^ cross section of the bone marrow. Trabecular bone is defined in the center of the grid and initially consists of 12.9% of the total area, consistent with our *in vivo* data (Supplementary Methods 1.1). Normal bone remodeling events are uniformly distributed over 4 years, the approximate turnover time of trabecular bone^83^. A remodeling event is initiated by the expression of RANKL by five osteoblast lineage cells on the perimeter of the bone.

#### Preosteoclasts and osteoclasts

Precursor osteoclasts follow the gradient of RANKL to the bone remodeling site. With a given probability, at least five preosteoclasts will fuse together to become a multinucleated osteoclast. Active osteoclasts resorb bone matrix, leading to the release of BDF. We assume that the amount of bone that osteoclasts resorb is proportional to their lifespan of approximately 14 days^83,98^.

#### MSCs, preosteoblasts, and osteoblasts

After an osteoclast fuses, an MSC is placed on the grid within a radius of 40 µm of the osteoclast, provided there is not already an MSC within the neighborhood. This requirement ensures that MSCs are located adjacent to sites of bone remodeling and can couple bone resorption with bone formation^36^. MSCs undergo asymmetrical division to create preosteoblasts, which proliferate when BDF is above a certain threshold or differentiate into bone matrix-producing osteoblasts when BDF is below the threshold (Supplementary Fig. 2c). We assume that osteoblast lifespan is proportional to the amount of bone that was resorbed at the location of the osteoblast, which is approximately 3 months. In the model, we assume that osteoblast death is proportional to the amount of bone that was resorbed by the osteoclast subunit that had occupied the space. Whilst this is a simplifying assumption of the biology, this assumption permits the coupling of bone resorption with bone formation, since the lifespan of an osteoblast controls how much bone the osteoblast forms. Bone resorption and bone formation are tightly coupled processes^99^. The definitive mechanisms through which osteoclasts and osteoblasts sense each other and their catabolic/anabolic processes have still not been fully elucidated and can be context dependent. However, several mechanisms are known to play a role including the release of bone derived factors (BDFs; namely TGF-beta and IGF-1), osteoclast derived factors such as sphingosine 1-phosphate, extracellular vesicles, apoptotic bodies containing RANK, and demineralized collagen remnants in the resorption lacunae. To model each of these mechanisms independently would make our model very complex as well as computationally expensive. We therefore integrated a straightforward coupling of osteoblasts and osteoclasts in the model. Further, when an osteoblast is buried, this has consequences for bone homeostasis. To maintain a steady state of bone, the lifespan of the nearest osteoblast is increased to allow it to build more bone to compensate for the amount lost due to the buried osteoblast. Whilst this is a simplifying assumption, research suggests osteoblast lineage cells communicate through gap junctions. This allows osteoblasts to coordinate activities and ensure new bone tissue is deposited at the correct location and in the appropriate amount to maintain structural integrity and bone homeostasis^100–104^.

#### Cytokines

RANKL (*R*_*L*_) and BDF (*T*_*β*_) are two key cytokines that drive the normal bone remodeling process. RANKL is produced by osteoblast lineage cells (*⍺*_*R*_*B_i,j_*) with natural decay of the ligand (*δ*_*R*_*R*_*L*_). BDF is released during bone resorption (*⍺*_*T*_*B_i,j_C_i,j_*) with natural decay of the ligand (*δ*_*T*_*T*_*β*_). We also assume that there is a constant production of BDFs (*⍺*_*B*_) by other cell types that are not explicitly defined in the model. These cytokines are incorporated into our model through the following system of partial differential equations (PDEs):

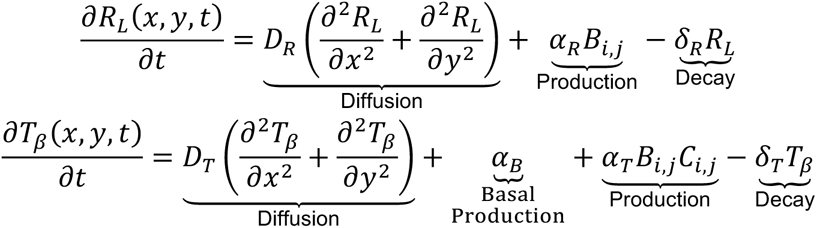

where the subscripts specify the location on the grid, i.e., *x* = *ih*, *y* = *jh* where {*i*, *j*} are positive integers and *h* denotes the spatial step, *h* = Δ*x* = Δ*y*. The system of PDEs is solved using the forward time centered space (FTCS) scheme with periodic boundary conditions imposed on all sides of the domain, which is implemented using the diffusion function in HAL.

#### Multiple Myeloma

A single myeloma cell is recruited within a radius of 40 *μ*m of an osteoclast after the first osteoclast fusion event occurs (Fig. 4a). MM cells move and proliferate in response to BDF and have a further proliferative advantage if they are within a radius of 20 *μ*m of an MSC or preosteoblast (Fig. 3g), and a survival advantage if BDF is above a certain threshold (Fig. 3h). As the MM cell population increases, additional MSCs are recruited. Furthermore, if three or more myeloma cells are located close to bone, a new bone remodeling event is initiated which increases RANKL and osteoclast formation. MM cells also prevent the differentiation of preosteoblasts if they are located within a radius of 80 *μ*m of a MM cell which decreases osteoblast activity.

#### Treatment

The HCA incorporates the direct and indirect effects of bortezomib by including the cytotoxic effect on MM cells as well as its ability to inhibit preosteoclast fusion and enhance bone formation^48,49^ (Supplementary Fig. 3). To mimic the clinical scenario of MM diagnosis, once MM reaches 10% of the bone marrow area *in silico*, treatment is applied either continuously (as for mouse models of multiple myeloma) or pulsed (2 weeks on, 1 week off; as is performed in the clinic) until MM burden reaches 20% of the bone marrow. In these simulations, once treatment is initiated, myeloma cells have a probability of developing resistance mechanisms during cell division that would cause the MM cells to become resistant to bortezomib. MM cells that are within a radius of 20 *μ*m of an MSC or preosteoblast or are located at a position where BDF is above a certain threshold, have a survival advantage during treatment referred to as environment mediated drug resistance (EMDR).

### Multiple myeloma mouse models

All animal experiments were performed with University of South Florida (Tampa, FL) Institutional Animal Care and Use Committee approval (CCL; #7356R). Male and female 14–16-week-old immunodeficient mice NOD-SCIDγ (NSG) mice were divided into tumor naïve or tumor bearing mice GFP-expressing human U266 (U266-GFP) multiple myeloma cells were injected (5×10^6^cells/100µL PBS) or PBS (100µL) via tail vein. Mice were euthanized at days 3, 10, 20, 40 and 100 (3-5/group/timepoint). In a separate experiment male and female 8 week-old NSG mice were inoculated with PI-sensitive U266-GFP-Luc (90%) and PI-resistant PSR-RFP (10%) MM cells via tail vein injection (5×10^6^cells/100µL PBS). Upon detection of tumor burden in the hind limbs via bioluminescent imaging (IVIS), mice were randomized to two groups to receive either twice weekly vehicle (PBS) or Zol (30µg/kg). Following one week of Zol pretreatment, mice were further randomized in to four subgroups and received the following: Group 1 Vehicle (5 mice), Group 2 Zol (30µg/kg/twice weekly; 5 mice), Group 3 Zol+BTZ (7 mice), Group 4 BTZ (0.5mg/kg/twice weekly; 4 mice). After one week of BTZ treatment, all mice were euthanized. Tibiae were excised and soft tissue removed for histological and FACs analyses.

### Micro-computed Tomography, Immunofluorescence and Histomorphometry

Harvested right tibiae were fixed in 4% paraformaldehyde (PFA) in 1x phosphate buffered saline (PBS) for 48 hours at room temperature. Evaluation of trabecular bone microarchitecture was performed in a region of 1000 µm, beginning 500μm from the growth plate using the SCANCO μ35CT scanner. Tibiae were decalcified in an excess of 10% EDTA at pH 7.4 at 4°C for up to three weeks with changes every 3 days before being place in cryoprotection buffer (30% sucrose), frozen in cryomedium (OCT) and subjected to cryosectioning to yield 20 μm tissue sections. Sections were dried, washed and blocked before addition of primary antibodies (anti-pHH3; 06-570 Millipore, anti-Osterix; ab22552, abcam: anti-αSMA: PA5-16697, Invitrogen). Slides were washed and incubated with goat anti-rabbit Alexa Fluor-647-conjugated secondary antibody and counterstained with DAPI. Mounted sections were imaged using 10X image tile-scans of whole tibia. pHH3+, αSMA+, Osterix+ or GFP+ cells were identified and quantified using Fiji software^105^. Dual ALP and TRAcP staining were performed on cryosections. Brightfield visualization was performed using the EVOS auto at 20X magnification. Five images per section were taken and the number of ALP+ cuboidal bone-lining osteoblasts and red multinucleated TRAcP+ osteoclasts per bone surface were calculated.

*For additional and in-depth methods, please see supplementary materials*.

## Author Contributions

A.K.M and D.B. were responsible for computational model design and execution. R.T.B and C.C.L were responsible for biological experimentation design and analysis. T.L, N.N, J.F performed biological experiments. P.R.S, R.C, A.S and K.H.S were involved in conception, design, experimentation, and analysis of *ex vivo* samples. A.K.M, R.T.B, C.C.L and D.B were involved in overall concept and design. All authors were involved in data interpretation, manuscript writing and editing.

## Acknowledgements

The authors would like to thank the patients at H. Lee Moffitt Cancer Center who provided clinical samples for *ex vivo* assays as well as consented to access to their clinical data through the Total Cancer Care database. In addition, the authors thank, Marilena Tauro, Tatiana Miti, Malgorzata Tyczynska, Mark Robertson-Tessi and Etienne Baratchart, at Moffitt Cancer Center for insightful discussions over the years and with editing and revision of manuscript. This work has been supported in part by the Small Animal Imaging Laboratory (SAIL) and the Flow Cytometry, and Analytical Microscopy Core Facilities at the Moffitt Cancer Center, an NCI designated Comprehensive Cancer Center (P30-CA076292). This work was supported in part by funds from the NCI PSON (5U01CA244101), Physical Sciences in Oncology (PSOC) Grant 1U54CA193489-01A1 (to K.H.S., A.S, and others), H. Lee Moffitt Cancer Center’s Team Science Grant (A.S., K.H.S.), Miles for Moffitt Foundation (A.S.). Additionally, this work was supported by personal donations from The Pentecost Myeloma Research Center.

## Data availability statement

All the data supporting the findings of this study are available within the article and its Supplementary information files and from the corresponding author upon reasonable request. A reporting summary for this article is available as a Supplementary Information file. Source data are provided with this paper.

## Code availability statement

The code generated in this manuscript is available at https://github.com/dbasanta/MM_ABM

## Supplementary Methods

### 1. Hybrid Cellular Automaton Methods

We developed a new hybrid cellular automaton (HCA) model to capture some of the key cellular and molecular populations that characterize the MM bone ecosystem. This model was developed from scratch although some of the assumptions, including those about what are the most important populations for the model, build on our previous experience building a similar model to study bone metastatic prostate cancer^1–3^. In this instance our code (available on this repository: https://github.com/dbasanta/MM_ABM) build on the HAL library which will help our code become a more self-sufficient and reusable platform of use to other bone-modelling scientists. We have generated different and novel data and *in vivo* model to parameterize, calibrate the model and validate the results. Building on our experience starting computational models of cancer from homeostasis, this new code is more robust and capable of performing bone remodeling simultaneously in several sites. This robustness is also demonstrated by the fact that homeostasis in the bone is preserved even with the use of variable depths of osteoclast-mediated bone resorption which was not the case previously.

The model we developed incorporates six different cell types, including precursor osteoclasts, active osteoclasts, mesenchymal stem cells (MSCs), precursor osteoblasts adult osteoblasts, and multiple myeloma (MM) cells. Additionally, we incorporate two signaling molecules including RANKL and bone derived factors (BDFs) such as transforming growth factor beta (TGF-β). In the sections below, we describe the specific assumptions and parameters incorporated into the HCA of normal bone remodeling (**Supplementary Section 1.1**), multiple myeloma (**Supplementary Section 1.2**), and bortezomib treatment (**Supplementary Section 1.3**). Refer to **Supplementary Tables 1-6** in **Supplementary Section 1.5** for a complete description of the parameters and values used in the HCA.

#### 1.1. Normal Bone Remodeling

##### Algorithmic Implementation

###### Initial and boundary conditions of agent grid

We first develop a model of trabecular bone remodeling in which we represent a cross section of the bone marrow as a 2D rectangular grid. We define the length of the grid as 0 ≤ *x* ≤ 160 *px* and the height as 0 ≤ *y* ≤ 150 *px*, where Δ*x* = 1 *px* = 10 *μm*, the diameter of a myeloma cell. To obtain the initial arrangement of bone, we first initiated the bone as a 60 *px* × 50 *px* rectangle in the center of the grid, surrounded by bone marrow. The initial area of the bone was set to represent the approximate bone area to total area (BA/TA) from sham limbs of wild type RAG2 mice (mean: 12.4%, n = 6, data not shown). As bone remodeling occurs over time, the initial rectangle is transformed, though the BA/TA remains approximately constant. We arbitrarily select one of these final bone configurations as the initial condition for subsequent runs of the model. Hence, trabecular bone initially consists of 12.9% of the total area. We assume that the number of precursor osteoclasts remains constant and is approximately equal to the number of monocytes, which is 2.8% of the bone marrow^4^. MSCs initially consist of 0.01% of the bone marrow^5^. We implement periodic boundary conditions on all sides of the domain for all cell types.

###### Timesteps

We use two separate time step sizes, Δ*t_*d*iff_* and Δ*t*_*cells*_, to account for the differences in the timescales for diffusion (fast) versus cell processes (slow). We use Δ*t_*d*iff_* = 1 *sec* and Δ*t*_*cells*_ = 6 *min*.

###### Simulation Process

The computational algorithm was implemented using the Hybrid Automata Library (HAL)^6^ and is organized as shown in **Supplementary Fig. 1a** and **Fig. 1c**.

##### Precursor osteoclasts

###### Fusion

Precursor osteoclasts fuse and become multinucleated osteoclasts. This process is regulated by the expression of RANKL (**Supplementary Fig. 2f**), which is expressed by osteoblast lineage cells such as osteoblast precursors, osteocytes, and bone lining cells^7,8^. It is not well understood which cell type is the primary source of RANKL for osteoclast fusion, but it has been suggested that microdamage to trabecular bone leads to osteocyte apoptosis, which may signal to bone lining cells to express RANKL^7,8^. In our model, we assume that bone remodeling events are initiated through microfractures that are uniformly distributed over 4 years, the approximate turnover time of trabecular bone^9^. The total number of remodeling events was approximated using the initial total amount of bone divided by the approximate amount of bone resorbed per osteoclast. For simplicity, each basic multicellular unit (BMU) initiates from a single remodeling event that consists of five adjacent cells on the perimeter of bone that begin to express RANKL. The number of cells expressing RANKL was selected to equal the approximate diameter of an osteoclast, which is 50 *μm*^10^.Precursor osteoclasts follow the gradient of RANKL and once there are five adjacent cells on the perimeter of bone, they fuse to become an active osteoclast (osteoclast) with probability

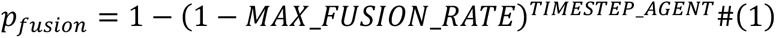

This probability reflects that fusion is a process that takes at least 3 days to complete. We note that this probability is scaled geometrically since the agent time step (6 min) is smaller than the unit of time (1 hour), hence the probability that the preosteoclast does not fuse per agent time step must be considered. Additionally, fusion requires stimulation with RANKL which is modeled as a Hill function such that the probability of fusion increases and saturates at one as RANKL increases (**Supplementary Fig. 2g**):

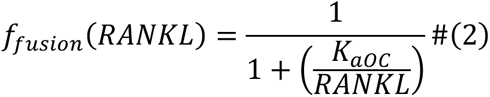

A Hill function is chosen as the functional form because it is commonly used to represent cell responses that are dependent on receptor-ligand interactions^11^. After the preosteoclasts fuse to become an osteoclast, five new preosteoclasts are randomly placed on the grid so that the number of preosteoclasts remains constant over time. To assess whether these functional forms qualitatively capture the dynamics of fusion in response to different concentrations of RANKL, we inhibit RANKL production by multiplying the production rate by scale factor *π*_*Ri*_(**Equation 10**). As expected, the total number of osteoclasts decreases as RANKL decreases (**Supplementary Fig. 2h**).

###### Movement

Precursor osteoclasts migrate in response to a gradient of RANKL^12,13^. We use the same technique to describe the probability of a cell remaining stationary or moving up, down, left, or right used in the original implementation of the hybrid discrete-continuum model^14^ (**Equation 9**). If a preosteoclast would otherwise move in a direction that is occupied by a myeloma cell or precursor osteoblast, the two cells swap positions on the grid. This requirement underlies the assumption that fusion is not limited by spatial restrictions and is necessary in order to capture *in vivo* data showing that osteoclasts increase over time due to multiple myeloma.

##### Active osteoclasts

In the model, active osteoclasts consist of 5 subunits to represent the size difference between osteoclasts and other cell types and to account for the fact that they are the result of the fusion of 5 preosteoclasts. In order for the osteoclast to function as a single unit, once a subunit of an osteoclast is selected for a given timestep, all of the other subunits are collected and the action for each is defined to be the same for a given osteoclast.

###### osteoclast lifespan

We assume that the amount of bone that osteoclasts resorb is proportional to their lifespan, which we define to be normally distributed with a mean lifespan of 14 days^9^, bounded between 7 and 21 days:

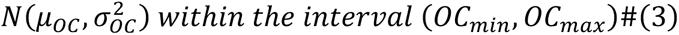

To verify the implementation of this assumption in the model, we show that the average osteoclast lifespan per year remains constant over time (**Supplementary Fig. 1e**). The end of the bone resorption phase is followed by a reversal period, which couples bone resorption and bone formation. When an osteoclast dies, it is removed from the grid, and the cleared space is designated as eroded bone. Bone derived factors continue to be produced at a lower level for three days, which allows precursor osteoblasts to move and attach to the eroded surface. This release of BDFs could either be due to matrix metalloproteinase activity^15^ or small osteoclasts that remain on the surface^16^, though the source is not modeled explicitly.

###### Osteoclast resorption

We assume that each osteoclast resorbs bone in the direction with the maximum amount of bone. Osteoclasts resorb bone tissue in areas where there is a high degree of mechanical stress or strain. There are several proposed mechanisms governing how osteoclasts detect areas of bone resorption. Integrins may play a role in guiding the direction of bone resorption by regulating the formation and orientation of the sealing zone. Specifically, integrins on the leading edge of the osteoclast are thought to sense the mechanical properties of the bone tissue and orient the osteoclast towards areas of higher mechanical stress or strain^17^. This results in the formation of a sealing zone that is oriented in the direction of the mechanical stress or strain, allowing the osteoclast to efficiently remove bone tissue from areas that are under the greatest mechanical load. Another hypothesis is that osteoclasts may be guided by the orientation of collagen fibers in bone^18^. Collagen is the main structural protein in bone, and it forms a network of fibers that give bone its strength and resilience. Studies have suggested that osteoclasts may be able to sense the orientation of collagen fibers in bone through a range of receptors and signaling pathways, and that this information may guide their resorption activity. Another hypothesis is that osteoclasts may be guided by chemical gradients of signaling molecules that are produced by other cells in the bone microenvironment^17^. By following these chemical gradients, osteoclasts may be able to target areas of bone where remodeling is needed.

To determine the direction, we check the Von Neumann neighborhood around each subunit of the osteoclast and record the total amount of bone in each direction. If more than one direction contains the maximum amount of bone, one of these directions is randomly selected. The bone is removed from the grid as it is resorbed and is replaced by the osteoclast. As bone is resorbed, bone derived factors are released from the bone matrix^16^, which is modeled as a production term in **Equation 10**. Because the amount of bone that osteoclasts resorb is proportional to their lifespan, we verify that the average amount of bone resorbed per osteoclast per year remains constant over time (**Supplementary Fig. 1g**).

##### Mesenchymal stem cells (MSCs)

###### Recruitment

MSCs have been identified in various niches within the bone marrow, and their precise location and distribution are still under investigation. Some of the most common locations where MSCs are found in the bone marrow are the endosteal region, the perivascular region, and the central marrow region^19,20^) MSCs are located in close proximity to the endosteal surface, where bone remodeling takes place, and the majority of MSCs are in contact with blood vessels or surrounded by pericytes and other undefined cells, which suggests that these cells are located in the perivascular niche^19,21^. After an osteoclast fuses, an MSC is placed on the grid within a certain radius of the osteoclast (MSC_radius), provided there is not already an MSC within the neighborhood. This requirement ensures that MSCs are located adjacent to sites of bone remodeling to couple bone resorption with bone formation^22^. However, to preserve that approximately 0.01% of the bone marrow consists of MSCs^5^, MSCs that are not within a certain radius of an osteoclast are removed from the grid. Therefore, the number of MSCs fluctuates over time but is capable of returning to the initial condition.

###### Proliferation

We assume that MSCs divide only when BDF is above a certain threshold (*BDF*_*t*O*res*O_) (**Supplementary Fig. 2c**). The probability of cell division is

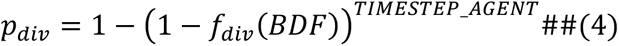

where

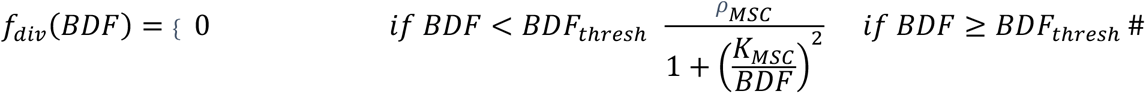

When an MSC divides we assume it undergoes asymmetric division, producing a precursor osteoblast. The daughter cell is randomly placed in an empty grid cell within the Moore neighborhood of the MSC.

###### Movement

MSCs migrate in the direction of higher TGF-β when there is a cytokine gradient, otherwise they move randomly^16^. As before, we use the hybrid discrete-continuum technique to describe the probability of cell movement (**Equation 9**).

##### Preosteoblasts

###### Death

We assume that preosteoblasts die if they are not adjacent to eroded bone with probability

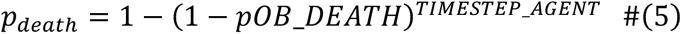

or with probability one if *pOB_*a*g*e*_* ≥ 42 *days*. Once a preosteoblast dies it is immediately removed from the grid.

###### Proliferation

We assume that high levels of BDF promote proliferation of preosteoblasts whereas low levels promote differentiation ^23^. The probability of cell division is the same as it is for MSCs (**Equation 4**; **Supplementary Fig. 2c**), except that a preosteoblast divides symmetrically to produce another preosteoblast. As before, the daughter cell is randomly placed in an empty grid cell within the Moore neighborhood of the preosteoblast. To assess whether these model assumptions qualitatively capture the dynamics of proliferation in response to different concentrations of BDF, we inhibit BDF production by multiplying the production rate by scale factor *π*_*Ti*_ (**Equation 10**). As expected, the average proportion of preosteoblast over time decreases as BDF decreases (**Supplementary Fig. 2e**).

###### Differentiation

After the bone resorption phase, preosteoblasts attach to the eroded surface and differentiate due to differentiation signals provided by the exposed bone matrix^22^. In the model, preosteoblasts that are adjacent to eroded bone and exposed to low levels of bone derived factors for two weeks will differentiate into active osteoblasts (osteoblasts).

###### Movement

We assume the precursor osteoblasts that are not attached to eroded bone, move in response to a gradient of bone derived factors, which is modeled in the same way as for MSCs with the hybrid discrete-continuum technique (**Equation 9**).

##### Osteoblasts

###### Death

Active osteoblasts ultimately either undergo apoptosis, become osteocytes, or become quiescent bone-lining cells^9^. In the model, we assume that osteoblast death is proportional to the amount of bone that was resorbed by the osteoclast subunit that had occupied the space prior to the osteoblast (osteoclast_depth). Therefore, osteoblast death is also proportional to the average time it takes for an osteoblast to form one unit of bone (basal_time). This assumption permits the coupling of bone resorption with bone formation, since the lifespan of an osteoblast controls how much bone the osteoblast forms. Thus, bone homeostasis is maintained even when the amount of bone resorbed by an osteoclast varies. Under normal conditions the lifespan of an osteoblast is approximately 3 months^9^, which we reproduce in our model (**Supplementary Fig. 1f**). This equation describes how to define the lifespan of an aOB so that it lives long enough to replace the bone resorbed by an aOC (during homeostasis).

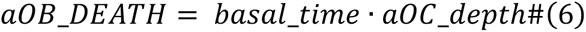

Units for equation 6:

◦ aOB_Death: days
◦ basal_time: days per unit of bone
◦ aOC_depth: units of bone

Alternatively, an active osteoblast may unintentionally get buried in bone by itself or other osteoblasts during bone formation. This is comparable to an osteoblast becoming an osteocyte, one of its potential cell fates. In the model, this occurs if the Von Neumann neighborhood of an osteoblast only contains bone or other osteoblasts. When an osteoblast is buried, this has possible consequences for bone homeostasis since the osteoblast did not form as much bone as it would have had it completed its full lifespan. To maintain a steady state of bone, the lifespan of the nearest osteoblast is increased to allow it to build more bone to compensate for the amount lost due to the buried osteoblast. This is a simplifying assumption of the model since biologically there are many factors involved to regulate the number and lifespan of osteoblasts. However, several studies have demonstrated the ability of osteoblasts, osteocytes and other cells of the BMU to sense and communicate with one another through juxtacrine and paracrine signaling to maintain bone homeostasis, through regulate cell proliferation, lifespan and activity ^24–27^.

###### Bone Formation

We assume that the time it takes for an osteoblast to form a unit of bone is dependent on the local concentration of BDF. To determine the functional relationship between BDF and mineralization rate we use *in vitro* data from MC3T3-E1 cells cultured in osteoblastic media with various concentrations of TGF-β or the TGF-β inhibitor, 1D11 (**Supplementary Methods 2.4**). This data shows how the mineralization rate compares to the control, specifically that it increases as TGF-β decreases (**Supplementary Fig. 2a**). We approximate this trend using an exponential decay function and extract the parameters using nonlinear least squares regression (**Supplementary Fig. 2b**):

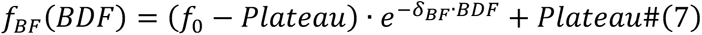

This equation is unitless since it represents fold change to bone mineralization time.

To determine the time that it takes an osteoblast to form one unit of bone as a function of BDF, we scale the time it takes an osteoblast to form one unit of bone in basal levels of BDF (basal_time) by **Equation 7**:

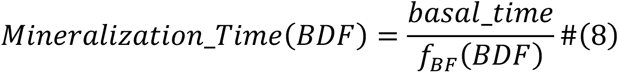

Units for equation 8:

◦ Mineralization_Time: days per unit of bone
◦ basal_time: days per unit of bone

This scaling allows for mineralization to increase when BDF is below the basal level. In the model, the mineralization time for each osteoblast is updated after sufficient time has passed to form a unit of bone. To determine which direction to form bone, we impose a set of rules to try to prevent the osteoblast from becoming buried and to prevent the creation of large gaps in the placement of the bone. To do this, we first check the Von Neumann neighborhood of an osteoblast for empty space. If more than one empty space exists, the osteoblast will preferentially move to the location where it will remain in contact with at least two units of bone and is not adjacent to another osteoblast; if this is not possible, one of the empty spaces will be randomly selected. To form bone, the osteoblast moves to the empty grid cell and its previous location is replaced with a new unit of bone. Because the amount of bone that osteoblasts produce is proportional to their lifespan, we verify that the average amount of bone produced per osteoblast per year remains constant over time (**Supplementary Fig. 1h**). To assess whether these model assumptions qualitatively capture the dynamics of mineralization in response to different concentrations of BDF, we inhibit BDF production by multiplying the production rate by scale factor *π*_*Ti*_ (**Equation 10**). As expected, BDF has a biphasic effect on bone in which low BDF results in bone loss due to lack of MSC/preosteoblast proliferation and medium BDF results in bone growth due to increased mineralization (**Supplementary Fig. 2d**).

##### Movement

In the model, MSCs, preosteoblasts, and MM cells migrate in the direction of higher TGF-β when there is a cytokine gradient, otherwise they move randomly^22^. preosteoclasts migrate in response to a gradient of RANKL^12,13^. We use the technique for motility described in the original implementation of the hybrid discrete-continuum model^14^ to describe the probability of each cell being stationary (*P*_0_), moving left (*P*_1_), right (*P*_2_), down (*P*_3_), or up (*P*_4_):

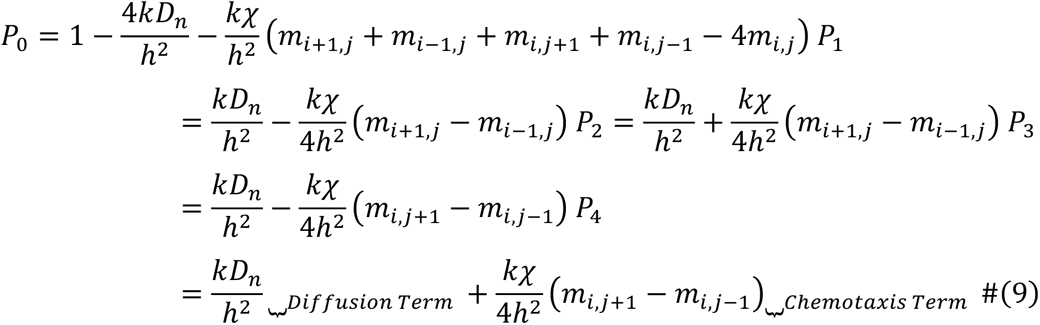

where the subscripts specify the location on the grid, i.e., *x* = *ih*, *y* = *jh* where {*i*, *j*} are positive integers, *k* denotes the timestep, *k* = *Δt*, and *h* denotes the spatial step, *h* = *Δx* = *Δy*. In the absence of a gradient of chemokine *m*, where *m* = {*R*_*L*_, *T*_*β*_}, cell movement is random with diffusion coefficient *D*_*n*_, whereas in the presence of a gradient cell movement is directed with chemotaxis coefficient *χ*.

##### Cytokines (RANKL and BDF)

Receptor activator of NF-kB ligand (RANKL; *R*_*L*_) is a cytokine that is produced in both soluble and membrane-bound forms and can activate osteoclasts. Recently, osteocytes have been identified as a major source of RANKL^28,29^. Once a microfracture occurs, RANKL is produced at the rate *⍺*_*R*_ at five adjacent cells on the perimeter of bone, which then diffuses in two spatial dimensions and degrades at the rate *δ*_*R*_.

Bone derived factors (BDFs; *T*_*β*_) consist of the group of cytokines that are released during bone remodeling, including TGF-β and IGF. These cytokines are produced in a latent form by many other cell types, including T-cells, macrophages, and platelets^30^. Instead of modeling the production of BDF explicitly by these other cell types, we assume that BDFs are constantly produced at rate *⍺*_*B*_, and further produced at rate *⍺*_*T*_ while bone is resorbed by osteoclasts. We assume that BDF also diffuses in two spatial dimensions and degrades at rate *δ*_*T*_.

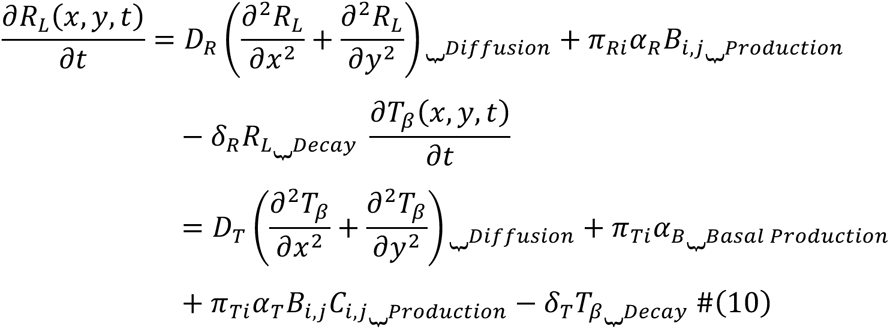

where

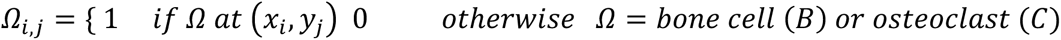

###### Boundary conditions of PDE grid

The system of PDEs is solved using the forward time centered space (FTCS) scheme with periodic boundary conditions imposed on all sides of the domain, which is implemented using the diffusion function in HAL. The timestep is chosen to ensure numerical stability, i.e., 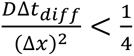.

#### 1.2. Incorporating Multiple Myeloma

##### Preosteoclasts

###### Fusion

Multiple myeloma cells enhance RANKL expression in the bone microenvironment which leads to increased osteoclastogenesis^31^. In the model, we assume that if there are three or more myeloma cells within the Moore neighborhood of a bone cell located on the perimeter of bone, a new bone remodeling event is initiated. As described above, each bone remodeling event is characterized by RANKL expression from five adjacent cells on the perimeter of bone.

##### Mesenchymal stem cells (MSCs)

###### Recruitment

Multiple myeloma cells produce chemoattractants that recruit MSCs to the bone marrow^32^. We assume that one MSC is recruited per 50 myeloma cells, and that once it is recruited, an MSC remains in the bone marrow (data not shown).

###### Proliferation

Myeloma cells inhibit the differentiation of mesenchymal stromal cells through soluble factors and cell-cell contract^33^. In the model, we assume that the differentiation of MSCs to precursor osteoblasts occurs through asymmetric division. To capture the effect of myeloma cells on osteoblast differentiation, we assume that MSCs do not divide if they are within a certain radius of a myeloma cell (MM_radius).

##### Preosteoblasts

###### Differentiation

Myeloma cells inhibit the differentiation of osteoblast progenitors through soluble factors and cell-cell contract^22^. In the model, we assume that preosteoblasts do not differentiate if they are within a certain radius of a myeloma cell (MM_radius),

##### Multiple Myeloma

###### Initiation

A single myeloma cell is recruited within a certain radius of an osteoclast after the first osteoclast fusion event occurs.

###### Death

The bone marrow microenvironment supports the growth and survival of multiple myeloma cells^34^. To determine where myeloma cell death and division occur spatially in the microenvironment, we measured the distance of these cells to the nearest bone (**Supplementary Methods 2.8**; **Fig. 3a-d**). Because there is less cell death close to bone, we assume that multiple myeloma cell death (sensitive and resistant cells) is dependent on the local concentration of BDF. The probability of cell death is

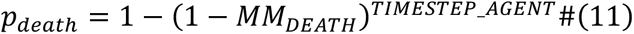

where

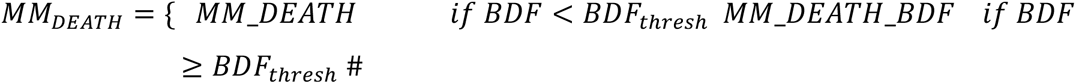

###### Proliferation

We assume that myeloma cell division is dependent on the concentration of BDF and has maximum rate *ρ*_*X*_. This maximum rate is dependent on if the myeloma cell is sensitive or resistant, since we showed that the U266 cell line has a cost of resistance (**Supplementary Fig. 7a-b**). Furthermore, we and others have shown that myeloma cell division increases near MSC/preosteoblast^35^ (**Fig. 3f** and **Fig. 6a**). To incorporate this into the model, we assume that sensitive cells within a certain radius of an MSC or preosteoblast divide faster than sensitive cells that are not close to an MSC or preosteoblast (**Fig. 3g**), which divide faster than resistant cells. We define the probability of cell division with the following Hill function:

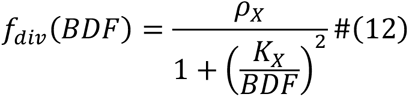

Where *ρ_X_* = {*MAX*_*MM*_*DIV*, *MAX*_*MM*_*DIV*_*MSC*, *MAX*_*R*_*MM*_*DIV*} and *K_X_* = {*K*_*MM*_, *K*_*MM*_*MSC*_, *K*_*R*_*MM*_} depending on if the myeloma cell is sensitive, resistant, and/or within a certain radius of an MSC or preosteoblast (protect_radius). A Hill function is chosen as the functional form because it is commonly used to represent cell responses that are dependent on receptor-ligand interactions^11^.

#### 1.3. Incorporating Bortezomib

##### Preosteoclasts

###### Fusion

Bortezomib inhibits osteoclast differentiation *in vitro* in a dose-dependent manner^25^. To incorporate this into the model we multiply the probability of fusion (**Equation 2**) by a repressive Hill function so that the probability decreases as dose increases (**Supplementary Fig. 3d**):

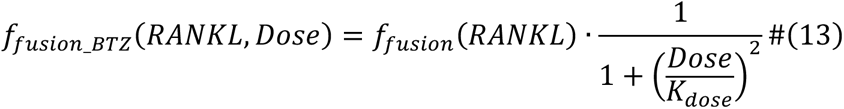

As expected, the total number of osteoclasts decreases as dose increases (**Supplementary Fig. 3g**).

##### Osteoblasts

###### Bone Formation

Bortezomib enhances the mineralization rate of osteoblasts *in vitro* in a dose-dependent manner (**Supplementary Fig. 3b**). To incorporate this into the model we scale the mineralization time (**Equation 8**) linearly so that at maximum dose (Dose = 1) the time it takes an osteoblast to form a unit of bone is reduced by half (**Supplementary Fig. 3c**):

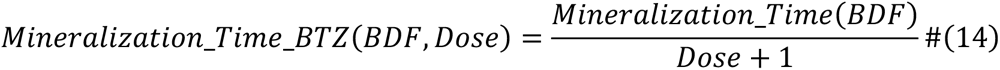

As expected, the BA/TA increases as dose increases (**Supplementary Fig. 3f**) due to increased production of bone per osteoblast.

##### Multiple Myeloma

###### Death

We assume that Bortezomib treatment does not affect the probability of cell death for resistant cells. On the other hand, Bortezomib decreases viability for sensitive cells *in vitro* and slows growth *in vivo* in a dose-dependent manner (**Supplementary Fig. 3a** and **Fig. 6c**)^36^. *In vivo*, failure to eradicate the disease may be due to both cell intrinsic mechanisms such as mutations, alterations in signaling pathways, copy-number alterations, epigenetic changes, and cell extrinsic mechanisms such as environment mediated drug resistance (EMDR)^37^. EMDR is when myeloma cells are transiently protected from Bortezomib due to factors or interactions with other cells in the bone microenvironment, which we define in our model to be when bone derived factors are above a certain threshold or MSCs or preosteoblasts are within a certain radius of a myeloma cell^32^. The probability of cell death during Bortezomib treatment is (**Supplementary Fig. 3i**):

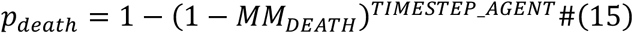

where

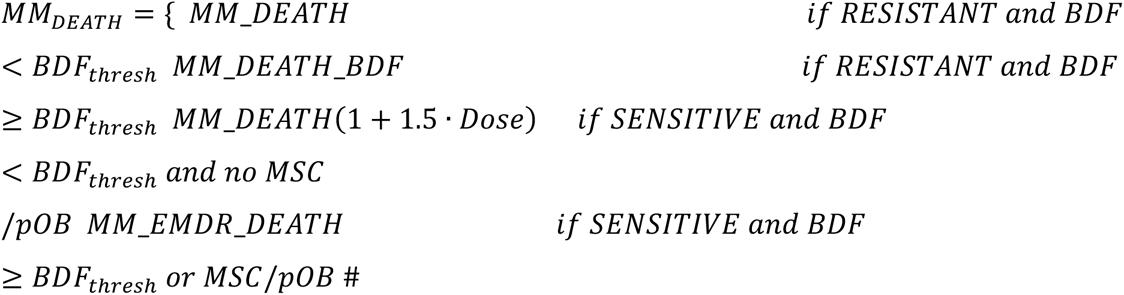

Therefore, tumor response to Bortezomib treatment depends on dose as well as the presence of bone derived factors or MSCs/preosteoblasts. At high dose, it is possible to completely eradicate the tumor in the absence of EMDR, similar to what is observed in *in vitro* experiments (**Supplementary Fig. 3j**).

###### Proliferation

Once treatment is initiated, myeloma cells have a probability of developing resistance (p_Ω_) during cell division that would cause the cell to become resistant to bortezomib.

##### Dose selection

###### Bortezomib

We define dose to be between 0 and 1, where 0 represents no treatment and 1 represents the maximum dose, i.e., the dose that is sufficient to kill off sensitive cells without having a major impact on normal bone cells such as MSCs and preosteoblasts. Based on the *in vitro* cell viability assay, we define maximum dose to be 10 nM (**Supplementary Fig. 3a**).

#### 1.4. Parameter Exploration

Parameter estimates used in the HCA model may vary widely based on the experimental design. For example, MM cell proliferation and death rates vary depending on the MM cell line used, growth media, and/or cell-cell interactions. To test whether our results are dependent on a particular parameter value, select parameters were varied to assess how the relapse time (when MM burden reached 20% of the marrow) and proportion of sensitive cells at endpoint changed under continuous bortezomib treatment (**Supplementary Fig. 10**). We found that the parameter controlling the impact of EMDR had a more striking effect on the proportion of sensitive cells at endpoint instead of the relapse time (**Supplementary Fig. 10b**), whereas the parameter controlling the cost of resistance had an impact on both outputs in the presence of EMDR (**Supplementary Fig. 10c**). Furthermore, the parameter controlling the survival advantage of BDF under no BTZ treatment conditions had only a minor impact on either output (**Supplementary Fig. 10d**), whereas the parameter controlling the proliferative advantage of sensitive cells in the presence of MSC/preosteoblast had a strong effect on the proportion of sensitive cells at endpoint. However, when the proliferative advantage is sufficiently strong, minimal residual disease may be present in the presence or absence of EMDR, though the proportion is higher with EMDR (**Supplementary Fig. 10e**). This analysis shows that although the proportion of sensitive cells at endpoint varied, our finding that EMDR maintains a reservoir of sensitive cells was robust for these parameters. Furthermore, we showed that EMDR consistently led to a higher proportion of tumors that relapsed, except when the proliferative advantage due to MSC/preosteoblast was sufficiently large.

#### 1.5. Parameters

The HCA model contains a large number of parameters as it has six different cell types and two microenvironmental factors. Many of the parameters were difficult to obtain and had to be estimated. Because many of the cell actions, such as proliferation and chemotaxis, result from the combination of parameters rather than a single value, our goal is to capture key outputs such as bone remodeling homeostasis instead of precisely parameterizing the model.

**Supplementary Table 1:** Grid Parameters.

| Grid Parameter | Description | Value | Units | Source |
| --- | --- | --- | --- | --- |
| *SPACESTEP*<br>($\Delta x$) | Space step of HCA grid | 10.0 | $\mu m$ | Estimated* |
| *TIMESTEP_AGENT*<br>($\Delta t_{cells}$) | Cell timestep | 0.1 | hour | Assumed |
| *N_TIMESTEP_PDE* | Number of diffusion timesteps per agent timestep | 360 | number | Assumed |
| *TURNOVER_TIME* | Turnover time of trabecular bone | 35040 | hours | <sup>9</sup> |
| *xDim* | Length of HCA grid | 1600 | $\mu m$ | Assumed |
| *yDim* | Height of HCA grid | 1500 | $\mu m$ | Assumed |

- *SPACESTEP*. We assume that 1 *pixel* = 10 *μm*, the diameter of a myeloma cell (experimentally derived)

**Supplementary Table 2:**
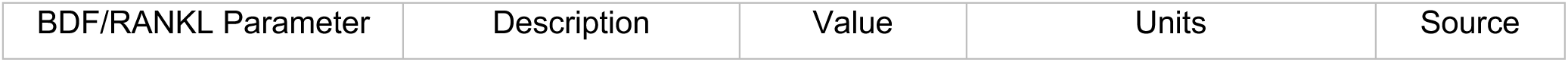
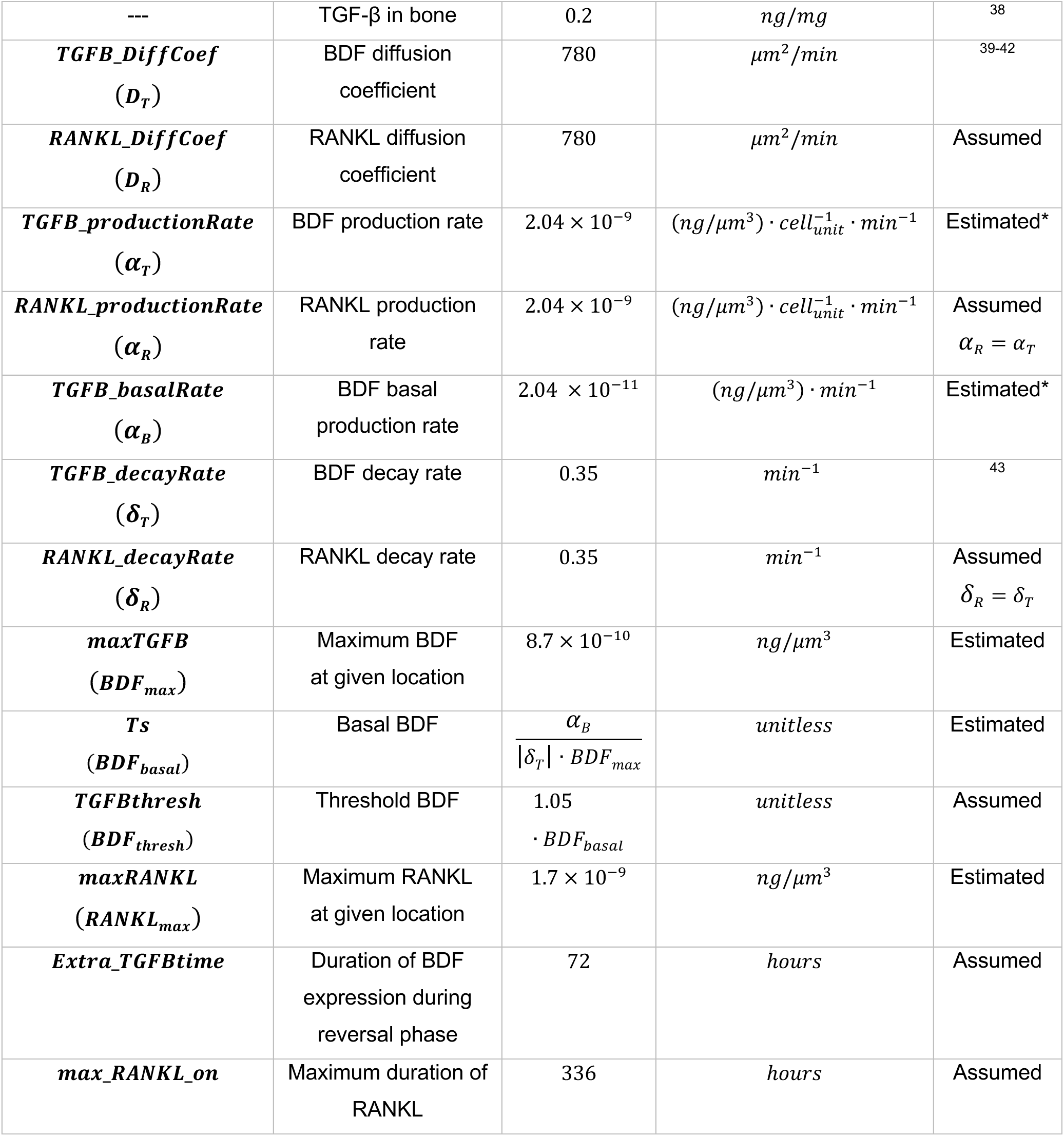
BDF/RANKL Parameters.

| BDF/RANKL Parameter | Description | Value | Units | Source |
| --- | --- | --- | --- | --- |
| --- | TGF-β in bone | 0.2 | $ng/mg$ | 38 |
| ***TGFB_DiffCoef***<br>($D_T$) | BDF diffusion coefficient | 780 | $\mu m^2/min$ | 39-42 |
| ***RANKL_DiffCoef***<br>($D_R$) | RANKL diffusion coefficient | 780 | $\mu m^2/min$ | Assumed |
| ***TGFB_productionRate***<br>($\alpha_T$) | BDF production rate | $2.04 \times 10^{-9}$ | $(ng/\mu m^3) \cdot cell_{unit}^{-1} \cdot min^{-1}$ | Estimated* |
| ***RANKL_productionRate***<br>($\alpha_R$) | RANKL production rate | $2.04 \times 10^{-9}$ | $(ng/\mu m^3) \cdot cell_{unit}^{-1} \cdot min^{-1}$ | Assumed<br>$\alpha_R = \alpha_T$ |
| ***TGFB_basalRate***<br>($\alpha_B$) | BDF basal production rate | $2.04 \times 10^{-11}$ | $(ng/\mu m^3) \cdot min^{-1}$ | Estimated* |
| ***TGFB_decayRate***<br>($\delta_T$) | BDF decay rate | 0.35 | $min^{-1}$ | 43 |
| ***RANKL_decayRate***<br>($\delta_R$) | RANKL decay rate | 0.35 | $min^{-1}$ | Assumed<br>$\delta_R = \delta_T$ |
| ***maxTGFB***<br>($BDF_{max}$) | Maximum BDF at given location | $8.7 \times 10^{-10}$ | $ng/\mu m^3$ | Estimated |
| ***Ts***<br>($BDF_{basal}$) | Basal BDF | $\frac{\alpha_B}{\vert \delta_T\vert \cdot BDF_{max}}$ | unitless | Estimated |
| ***TGFBthresh***<br>($BDF_{thresh}$) | Threshold BDF | $1.05 \cdot BDF_{basal}$ | unitless | Assumed |
| ***maxRANKL***<br>($RANKL_{max}$) | Maximum RANKL at given location | $1.7 \times 10^{-9}$ | $ng/\mu m^3$ | Estimated |
| ***Extra_TGFBtime*** | Duration of BDF expression during reversal phase | 72 | hours | Assumed |
| ***max_RANKL_on*** | Maximum duration of RANKL | 336 | hours | Assumed |

- *TGF-β diffusion rate, D*_*T*_. The 780 *μm*^2^/*min* was estimated using the molecular weight of TGF-β (25 kDa ^39^). A molecule of molecular weight (M) 0.3-0.5 kDa has a diffusion coefficient of approximately 10^−6^ *cm*^2^/*s* ^40^. The plot of *log log* (*D*) vs. *log log* (*M*) correlates with a line of slope 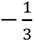 ^41^. Thus, we solve

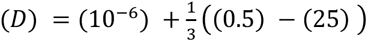 to get *D* = 2.6 × 10^−7^ *cm*^2^/*s* = 1560 *μm*^2^/*min*. Small molecules diffuse through the extracellular space with an effective diffusion coefficient that is two to three times less than the free diffusion coefficient^42^, thus *D*^∗^ = *D*/2 = 780 *μm*^2^/*min*.
- *TGF-β production rate, ⍺*_*B*_.

◦ Volume of resorption pit: *πr*^2^*h* = *π*(25^2^)(10) = 19634.95 *μm*^3^/*day*, where *r* = radius of osteoclast and *h* = amount of bone resorbed per day
◦ Rate of bone resorption by a 50 *μm* OC: Given the density of bone = 1500 *kg*/*m*^3 44^, (1500 *kg*/*m*^3^) × (19634.95 *μm*^3^/*day*) × (10^−18^ *m*^3^/*μm*^3^) = 2.945 × 10^−11^ *kg*/*day* = 2.945 × 10^−5^ *mg*/*day*
◦ TGF-β released/day due to bone resorption by a 50 *μm* OC: Given concentration of TGF-β in bone is 0.5 ng/mg ^38^, (2.945 × 10^−5^ *mg*/*day*) × (0.5 *ng*/*mg*) = 1.47 × 10^−5^ *ng*/ *day*. Because a 50 *μm* OC takes up 5 pixels, we divide this by 5 to get: 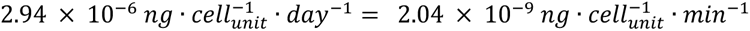, consistent with^45^. We convert this to units of ng per volume to be consistent with the units of chemotaxis, which requires dividing the amount (ng) by the soluble volume of each grid point. Because the soluble volume fraction is unknown, we set 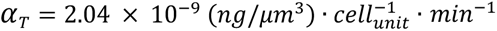, which was calibrated to reproduce the steps of normal bone remodeling.
- *TGF-β basal production rate, ⍺*_*B*_. We assume that BDF is produced at a higher rate through bone resorption compared to other sources in HCA^22^
- *TGF-β decay rate, δ*_*T*_ : The half-life of *TGF-β* is 2 minutes^44^. Thus, the decay rate is 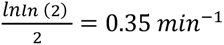.
- maxTGFꞵ was determined by running the unnormalized model a number of times and then recording the maximum TGFꞵ at a single grid point. This value of BDF was then set as maxTGFꞵ. The maximum TGFꞵ is printed after each model is run (tmax) so that it can be monitored whether the value stays between 0 and 1.
- Ts is the basal BDF and is defined as written in the table, *⍺*_*B*_/(|*δ*_*T*_| · *BDF*_*max*_). This is the amount of BDF that is always present on the grid.
- MaxRANKL was determined by running the unnormalized model a number of times and then looking at the output “preosteoblastborn.csv” to evaluate the maximum value of RANKL that occurred when an osteoclast formed. This value of RANKL was then set as maxRANKL so that most values of RANKL that lead to the formation of an osteoclast are between 0 and 1 (however, it is possible that some values in future runs of model are not strictly below 1).

**Supplementary Table 3:** preosteoclast/osteoclast Parameters.

| preosteoclast/osteoclast Parameter | Description | Value | Units | Source |
| --- | --- | --- | --- | --- |
| *pOC_DiffCoef* | preosteoclast random motility | 0.3 | $\mu m^2/min$ | Estimated |
| *pOC_TaxisCoef* | preosteoclast chemotaxis coefficient | $5 \times 10^{10}$ | $\mu m^2 min^{-1} (ng/\mu m^3)^{-1}$ | Estimated |
| *K<sub>aoc</sub>* | Half-maximal concentration constant | 0.01 | *concentration* | Assumed |
| *μ<sub>oc</sub>* | Mean osteoclast lifespan | 14 | *days* | 9,46 |
| *σ<sub>oc</sub>* | osteoclast lifespan standard deviation | 7/3 | *days* | Assumed |
| *OC<sub>min</sub>* | Minimum osteoclast lifespan | 7 | *days* | Assumed |
| *OC<sub>max</sub>* | Maximum osteoclast lifespan | 21 | *days* | Assumed |
| *MAX_FUSION_RATE* | Maximum fusion rate | 1/72 | *hours*<sup>-1</sup> | 47 |
| *Unit_RESORPTION* | Time to resorb one unit of bone | 1 | *day* | 48 |
| *aOC diameter* | Diameter of osteoclast | 50 | $\mu m$ | 10 |

- *Random motility coefficient of cell, preosteoclast_DiffCoef*: Stokes-Einstein relation: 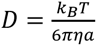. Given *k*_*B*_ = 1.38 × 10^−23^ *N* · *m*, *T* = 310 *K* (body temp: 310 *K*; room temp: 300 *K*), *η* = 10^−3^ *N* · *s*/*m*^2^ (viscosity of water), *a* = 5 *μm* (cell radius), *D* = 4.54 × 10^−10^*cm*^2^*s*^−1^ = 3 *μm*^2^/*min*. We assume that preosteoclast random motility is 10-fold slower in bone marrow compared to water.

**Supplementary Table 4:** MSC/preosteoblast/osteoblast Parameters.

| MSC/preosteoblast/osteoblast Parameter | Description | Value | Units | Source |
| --- | --- | --- | --- | --- |
| *MSC_DiffCoef*<br>*= pOB_DiffCoef* | MSC/preosteoblast random motility | 0.03 | $\mu m^2/min$ | Estimated |
| ***MSC_TaxisCoef*** | MSC<br>chemotaxis<br>coefficient | $5 \times 10^9$ | $\mu m^2 min^{-1} (ng/\mu m^3)^{-1}$ | Estimated |
| ***pOB_TaxisCoef*** | preosteoblast<br>chemotaxis<br>coefficient | $5 \times 10^{11}$ | $\mu m^2 min^{-1} (ng/\mu m^3)^{-1}$ | Estimated |
| ***K_{MSC} = K_{pOB}*** | Half-maximal<br>concentration<br>constant | $\sqrt{3} \cdot BDF_{thresh}$ | *unitless* | Assumed |
| ***MAX_MSC_DIVISION_RATE***<br>***= MAX_pOB_DIVISION_RATE***<br>***($\rho_{MSC} = \rho_{pOB}$)*** | Maximum<br>proliferation<br>rate | 1/24 | *hour*<sup>-1</sup> | Assumed |
| ***pOB_DEATH*** | preosteoblast<br>death rate | 1/72 | *hour*<sup>-1</sup> | Assumed* |
| ***pOB_DIFF*** | preosteoblast<br>differentiation<br>time | 336 | *hour* | Determined<br>from<br>experiments |
| ***MSC_radius*** | Radius<br>around<br>osteoclast/M<br>M to recruit<br>MSC | 40 | $\mu m$ | Assumed* |
| ***MM_radius*** | Radius of MM<br>effect on<br>MSC/preoste<br>oblast<br>differentiation | 80 | $\mu m$ | Assumed* |
| ***Y0***<br>***($f_0$)*** | Maximum fold<br>change | 5.055 | *unitless* | Estimated* |
| ***Plateau*** | Minimum fold<br>change | 0.5379 | *unitless* | Estimated* |
| ***scalefactor*** | Scale factor | $\frac{19.2}{BDF_{basal}}$ | *unitless* | Estimated* |
| ***k***<br>***($\delta_{BF}$)*** | Fold change<br>decay<br>constant | $0.1136 \cdot scalefactor$ | *unitless* | Estimated* |
| ***basal_time*** | Basal unit<br>bone<br>formation<br>time | 9 | *days* | 49 |

- *Random motility coefficient of cell, MSC_DiffCoef, preosteoblast_DiffCoef*: We assume that MSC and preosteoblast random motility is 100-fold slower in bone marrow compared to water, and 10 fold slower than preosteoclast.
- *Chemotaxis coefficient, MSC_TaxisCoef, preosteoblast_TaxisCoef*: The chemotaxis coefficient of human neutrophils in response to the tripeptide FNLLP is 150 *cm*^2^*s*^−1^*M*^−1^ ≈ 5.18 × 10^−2^*mm*^2^(*pg*/*mm*^3^)^−1^*day*^−1^ = 3.6 × 10^13^*μm*^2^(*ng*/*μm*^3^)^−1^*min*^−1^ (using 25 kDa as the molecular weight of TGF-β)^50,51^. However, to give reasonable model outputs required us to calibrate this parameter further and assume that MSC_TaxisCoef < preosteoblast_TaxisCoef.
- *preosteoblast death rate, preosteoblast_DEATH*: Assumed to be same as preosteoclast^47^
- *MSC recruitment radius, MSC_radius*: Defined so that MSC is close enough to respond to BDF generated by osteoclast
- *MM radius of impact on MSC/preosteoblast, MM_radius*: Defined so that MSC/preosteoblast responds to MM even when not directly adjacent. This assumption was necessary since HCA assumes MM is unable to migrate towards MSC/preosteoblast in absence of BDF, which created distance between MM and MSC/preosteoblast.
- *Exponential decay* (**Equation 7**)*: Y0, Plateau, k*: These values were estimated by using an exponential decay function and nonlinear least squares regression with *in vitro* data from MC3T3-E1 cells cultured in osteoblastic media with various concentrations of TGF-β or the TGF-β inhibitor, 1D11 (**Supplementary Methods 2.4**).

**Supplementary Table 5:** MM Parameters.

| MM Parameter | Description | Value | Units | Source |
| --- | --- | --- | --- | --- |
| ***MM_DiffCoef*** | MM random motility | 0 | $\mu\text{m}^2/\text{min}$ | Assumed |
| ***MM_TaxisCoef*** | MM chemotaxis coefficient | $5 \times 10^9$ | $\mu\text{m}^2 \text{ min}^{-1} (\text{ng}/\mu\text{m}^3)^{-1}$ | Estimated |
| ***MM_DEATH*** | MM basal death rate | 1/120 | $\text{hour}^{-1}$ | Assumed* |
| ***MM_EMDR_DEATH*** | MM death rate with EMDR | 1/120 | $hour^{-1}$ | Assumed* |
| ***MM_DEATH_BDF*** | MM death rate with BDF | 1/1200 | $hour^{-1}$ | Assumed* |
| ***MAX_MM_DIVISION_RATE*** | Maximum proliferation rate | 1/24 | $hour^{-1}$ | Estimated |
| ***MAX_MM_DIV_MSC*** | Maximum proliferation rate | 1/24 | $hour^{-1}$ | Assumed |
| ***MAX_RESISTANT_DIVISION_RATE*** | Maximum proliferation rate | 1/48 | $hour^{-1}$ | Estimated |
| ***K<sub>MM</sub>*** | Half-maximal concentration constant | $\sqrt{3} \cdot BDF_{thresh}$ | *unitless* | * |
| ***K<sub>MM_MSC</sub>*** | Half-maximal concentration constant | $BDF_{basal}$ | *unitless* | * |
| ***K<sub>R_MM</sub>*** | Half-maximal concentration constant | $\sqrt{1.25} \cdot BDF_{basal}$ | *unitless* | * |
| ***protect_radius*** | Radius of MM birth/death advantage from MSC/preosteoblast | 20 | $\mu m$ | Assumed |
| ***p<sub>resistance</sub>(p<sub>Ω</sub>)*** | Resistance probability | *variable* | *unitless* | Assumed |

- *Random motility coefficient of cell, MM_DiffCoef*: We assume that MM cells do not have random motility.
- *Chemotaxis coefficient, MM_TaxisCoef*: We assume that MM_TaxisCoef = MSC_TaxisCoef.
- *MM death rates, MM_DEATH, MM_EMDR_DEATH, MM_DEATH_BDF*: We assume that the lifespan of MM is similar to short-lived (3-5 days) and long-lived (several months) plasma cells^52^
- *Half maximal constant, K_MM_*. We set 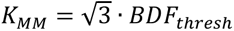, the value of BDF for which the myeloma proliferation rate is half maximum. This is estimated under the assumption that when *BDF* = *BDF_thresh_*, i.e., when BDF is slightly above the basal level that is constantly present, the proliferation rate = 1/96 hour^−1^. This value is based on the doubling time of myeloma cells *in vivo*.
- *Half maximal constant, K_MM_MSC_*. We set *K*_*MM*_*MSC*_ = *BDF*_*basal*_, which is estimated under the assumption that when *BDF* = *BDF*_*basal*_, i.e. in locations in the bone marrow that are not close to bone resorption, the myeloma proliferation rate = 1/48 hour^−1^ in the presence of MSC/preosteoblast (i.e. half the maximum proliferation rate).
- *Half maximal constant, K_R_MM_*. We set 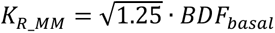, which is estimated so that when *BDF* = *BDF*_*basal*_, the division rate of resistant myeloma cells is less than the division rate of sensitive myeloma cells and remains greater than the death rate in the absence of Bortezomib treatment.

**Supplementary Table 6:** BTZ Parameters.

| BTZ Parameter | Description | Value | Units | Source |
| --- | --- | --- | --- | --- |
| $K_{dose}$ | Half-maximal concentration constant | 0.05 | *unitless* | Assumed |

### 2. Biological Supplementary Methods

#### 2.1. Cell culture

Human proteasome inhibitor sensitive (U266) myeloma cell and their proteasome inhibitor resistant derivatives (PSR^53,54^) were a kind gift from Dr. Steven Grant at the University of Virginia, VA. U266 and PSR were transduced using QIAGEN lentiviral particles (CLS-PCG-8 or CLS-PCR-8) according to manufacturer’s instructions to express GFP and RFP respectively, generating U266-GFP and PSR-RFP. These cells were cultured in RPMI containing 10% FBS (PEAK), 1% penicillin-streptomycin. MCSF-generated macrophages were isolated as previously described^55,56^. Briefly, tibia and femur were harvested from 6-week-old C57Bl/6 RAG2-/-mice. Bone marrow cells were collected by centrifugation (10,000 g, 15s) cells were plated aMEM (+/+) containing 10% FBS (Peak), 1% penicillin-streptomycin and MCSF (300-25, Peprotech; 30ng/ml). Adherent macrophages were collected and used for downstream experiments after 72 hours. Murine mesenchymal stromal cells (MSCs) were isolated as previously described^57^. Human MSCs (PT-2501) and the murine preosteoblast cell line, MC3T3-E1 were purchased from Lonza and ATCC, respectively. Mouse MSCs and MC3T3-E1 cells were cultured in aMEM (−/−) containing 1% penicillin-streptomycin and 15 or 10% FBS (PEAK), respectively. Human MSCs were cultured as above except with 10% qualified FBS (Gibco).

#### 2.2. MTT Assay

Myeloma cell lines, mouse MSCs or MC3T3-E1 were plated in 96-well plates at a density of 1×10^4^ cells/well. Cells were treated with vehicle or a range of concentrations of zoledronate or bortezomib. Cell viability was assessed at 72 hours by the MTT assay following the manufacturer’s instructions (CellTiter 96, #G3582, Pierce.) The absorbance was measured at 490nm after 3 hours of incubation at 37°C.

#### 2.3. Osteoclastogenesis Assay

Osteoclast formation assays were performed using murine MCSF-generated macrophages as previously described^55,56^. Briefly, MCSF-generated macrophages from 4–6-week-old C57Bl/6 RAG2-/- mice were plated in a 96-well plate (25×10^3^ cells/well) in triplicate with MCSF (30ng/ml) and allowed to adhere for 24 hours. RANKL (315-11C, Peprotech) was given at the indicated doses with M-CSF on days 1 and 3. Cultures were fixed with 4% paraformaldehyde on day 5 and TRAcP stained. TRAcP positive multinucleated (>3) were deemed osteoclasts.

#### 2.4. Osteoblast mineralization assay

Mouse MSCs, MC3T3-E1 cells or human MSCs (huMSCs) were differentiated with 1X StemXVivo mouse/rat or human osteogenic supplement (CCM009; R&D Systems) for indicated number of days. To assess the effect increased TGF-β has on OB differentiation, either exogenous human TGF-β1 (240-B; R&D systems) or vehicle (PBS) was added to cultures. To assess the effect decreased TGF-β levels have on OB differentiation, either anti-TGF-β antibody (1D11; R&D systems) or isotype control (13C4; R&D systems) was added to cultures. OB mineralization was assessed by Alizarin red staining as previously described^55,56^. Conditioned medium was generated from huMSCs or huMSCs differentiated for 7-28 days in osteogenic supplement by incubating cells in standard aMEM for 24hours.

#### 2.5. MM-MSC and sensitive/resistant MM co-culture assays

Human MSCs (huMSCs; Lonza) were plated (2000 cells/100ul) or media alone in a 96 well plate and allowed to adhere overnight. The following day 50% of the media was removed and replaced with media containing U266-GFP+ MM cells in standard RPMI. Plates were centrifuged at 200g for 5 minutes to facilitate adhesion to MSCs. After 3 hours, bortezomib was added at indicated concentrations. Images were taken at 0 and 72 hours. The area covered by GFP+ MM cells was calculated in Fiji software^58^. U266-GFP and PSR-RFP (10,000 total cells/well,) were plated at indicated ratios in a 96-well plate with indicated concentrations of bortezomib. Cell confluency for GFP and RFP populations was measured every 12 hours using Incucyte S3. For long term (60 day and 30 day cultures). Cultures were set up as previously described. BTZ (10nM) was given over 72 hour periods and PI-sensitive U266 and PI-resistant PSR confluency was measured by fluorescent markers using the incucyte SX5. After treatment, media was replaced, after PBS washing, with standard RPMI. Images were taken at indicated timepoints with EVOS FL auto prior to media changes.

#### 2.6. Flow cytometry

Left tibiae were used to assess tumor burden by GFP expression. Tibial ends were excised, whole bone marrow was isolated by centrifugation at 10,000g for 10 seconds. Red blood cells were lysed by RBC lysis buffer (R7757, Sigma-Aldrich) as per manufacturers guidelines. Bone marrow cells were subject to viability staining with Zombie Near-Infrared (NIR; 1:500; 423105, BioLegend). Appropriate compensation and fluorescence-minus-one (FMO) controls were generated in parallel either with aliquots of bone marrow cells or U266^GFP+/−^. Stained controls and samples were analyzed using BD Biosciences LSRII flow cytometer. The percentage of GFP-expressing cells was gated on singlet live bone marrow cells in FCS Express 7. To address the proliferative advantages provided by cells of the BME to MM cells, U266 MM cells were stained with CM-DIL (Invitrogen; V22888) according to the manufacturer’s instruction and incubated with 50% (v/v) control, MSC, preosteoblasts (Day 7 or 14) or OB (day 21 or 28) conditioned media for 7 days, with media changes every 3 days. An aliquot of untreated CM-DIL+ cells was stained with Zombie NIR as above and used for baseline time point. At the end of the experiment, MM cells were stained with Zombie NIR for live/dead discrimination. The MFI for the CM-DiI channel was calculated at baseline and after 72 hours on live single cell cells using appropriate FMO controls. FCS Express 7 was used to calculate the proliferative index of these cells.

#### 2.7. Microcomputed Tomography

Harvested right tibiae were fixed in 4% paraformaldehyde for 48 hours. Tibiae from mice from all time points were centralized and were subjected to micro-computed topography (μCT) scanning using SCANCO μ35 scanner to elucidate bone volume data at the proximal tibial metaphases. Individual bone scans were deidentified using numerical codes during, and reidentified post analyses in a blinded fashion. Evaluation of trabecular bone microarchitecture was performed in a region that consisting of 1000 μm, beginning 500μm from the growth plate. A three-dimensional cubical voxel model of bone was built, and calculations were made for relative bone volume per total volume (BV/TV), trabecular number (Tb.N), trabecular thickness (Tb.Th), trabecular spacing (Tb.Sp) and connectivity density (Conn.D).

#### 2.8. Immunofluorescence

Tibiae were decalcified in an excess of 10% EDTA for 21 days, refreshing EDTA every 48 hours before being placed in 30% sucrose (w/v) for 48 hours and subsequent cryosectioning. Tibia were mounted on Superfrost Plus slides (Fisher) with optimal cutting temperature (OCT) media and frozen on dry ice. Using a cryostat, 20 µm sections were made and stored at −80°C. For immunofluorescence, sections were kept on a slide warmer overnight at 56°C. Sections were rehydrated for 30 minutes at room temperature in PBS. Sections were blocked with 10% (v/v) normal goat serum and washed in PBS before staining addition of primary antibodies. Sections were incubated overnight at 4°C with primary rabbit antibodies at a dilution of 1:100 (anti-pHH3; 06-570 Millipore, anti-Osterix; ab22552, abcam: anti-aSMA: PA5-16697, Invitrogen,). Subsequently, slides were washed thrice in PBS and stained with goat anti-rabbit Alexa Fluor-647-conjugated secondary antibody (1:1000) at room temperature for 1h. DAPI (1ng/ml) was used as a nuclear counterstain. Mounted sections were imaged using 10X image tile-scans of whole tibia. The number of pHH3+ nuclei that co-localized with GFP+ cells were counted and the distance to nearest bone (tibia or cortical) was calculated. For MSC analyses, the percentage of total area in the trabecular region that was covered by αSMA+ cells were calculated on Fiji software. For preosteoblasts, the number of Osterix+ nuclei were counted and normalized to perimeter of trabecular bone for each mouse (Osx^+^ cells/mm).

#### 2.9. Histomorphometry TRAcP and ALP analyses

Additional tibia bone sections were cryosectioned and baked at 42°C overnight to improve tissue adhesion while retaining endogenous enzymatic activity. Tibial bone sections were rehydrated in PBS before incubation in napthol-AS-MX phosphate (Sigma-Aldrich; 855-20) and fast blue RR salt (Sigma-Aldrich; FB25-10CAP) solution for 30 minutes to identify ALP+ regions with a dark blue stain. After washing in PBS, the same sections were incubated in basic stock solution for 5 minutes at 37°C. Tibial sections were developed in pararosaniline dye and sodium nitrite at 37°C for 3 minutes to highlight red TRAcP+ cells. Sections were further counterstained with hematoxylin to visualize bone morphology. Slides were mounted and imaged using EVOS Auto brightfield at 20X magnification. Five images per section were taken and the number of ALP+ cuboidal bone-lining osteoblasts and red multinucleated TRAcP+ osteoclasts per bone surface were calculated.

#### 2.10 Ex vivo mathematical myeloma advisor (EMMA) platform: Pentecost Myeloma Research Center cohort

An *ex vivo* assay was used to quantify the chemosensitivity of primary MM cells^59–61^. Fresh BM aspirate cells were enriched for CD138^+^ expression using Miltenyi (Bergisch Gladbach, Germany) 130-051-301 antibody-conjugated magnetic beads. MM cells (CD138^+^) were seeded in Corning (Corning, NY) CellBIND 384 well plates with collagen I and previously established human-derived stroma, containing approximately 4000 MM cells and 1000 stromal cells. Each well was filled with 80 μL of Roswell Park Memorial Institute (RPMI) 1640 media supplemented with fetal bovine serum (FBS, heat inactivated), penicillin/streptomycin, and patient-derived plasma (10%, freshly obtained from patient’s own aspirate, filtered) and left overnight for adhesion of stroma. The next day, drugs were added using a robotic plate handler so that every drug/combination was tested at 5 (fixed concentration ratio, for combinations) concentrations (1:3 serial dilution) in two replicates. Negative controls (supplemented growth media with and without the vehicle control dimethyl sulfoxide [DMSO]) were included, as well as positive controls for each drug (cell line MM1.S at highest drug concentration). Plates were placed in a motorized stage microscope (EVOS Auto FL, Life Technologies, Carlsbad, CA) equipped with an incubator and maintained at 5% CO_2_ and 37 °C. Each well was imaged every 30 min for a total duration of up to 6 days. A digital image analysis algorithm^59–61^ was implemented to determine changes in viability of each well longitudinally across the 96 or 144hr intervals. This algorithm computes differences in sequential images and identifies live cells with continuous membrane deformations resulting from their interaction with the surrounding extracellular matrix. These interactions cease upon cell death. By applying this operation to all 288 images acquired for each well, we quantified non-destructively, and without the need to separate the stroma and MM, the effect of drugs as a function of concentration and exposure time. Digital image analysis computes percent viability of MM cells for each time point and experimental condition (drug and concentration). For each patient-drug, we have a dose-time-response surface, which is abstracted into AUC (area under the curve), which is an area/integral measure of *ex vivo* response to therapy computed by taking an average of all *ex vivo* responses across all time (first 96h) and concentration. Clinical data was matched to patient samples ex vivo. Samples in the lowest AUC quartile were considered sensitive.

#### 2.11. Statistical Analysis

Statistical analysis was performed using the analysis of variance (ANOVA) with the appropriate post multiple comparison analysis in GraphPad prism 8.0-10.0.

### Supplementary Figures and Legends

**Supplementary Fig. 1.**
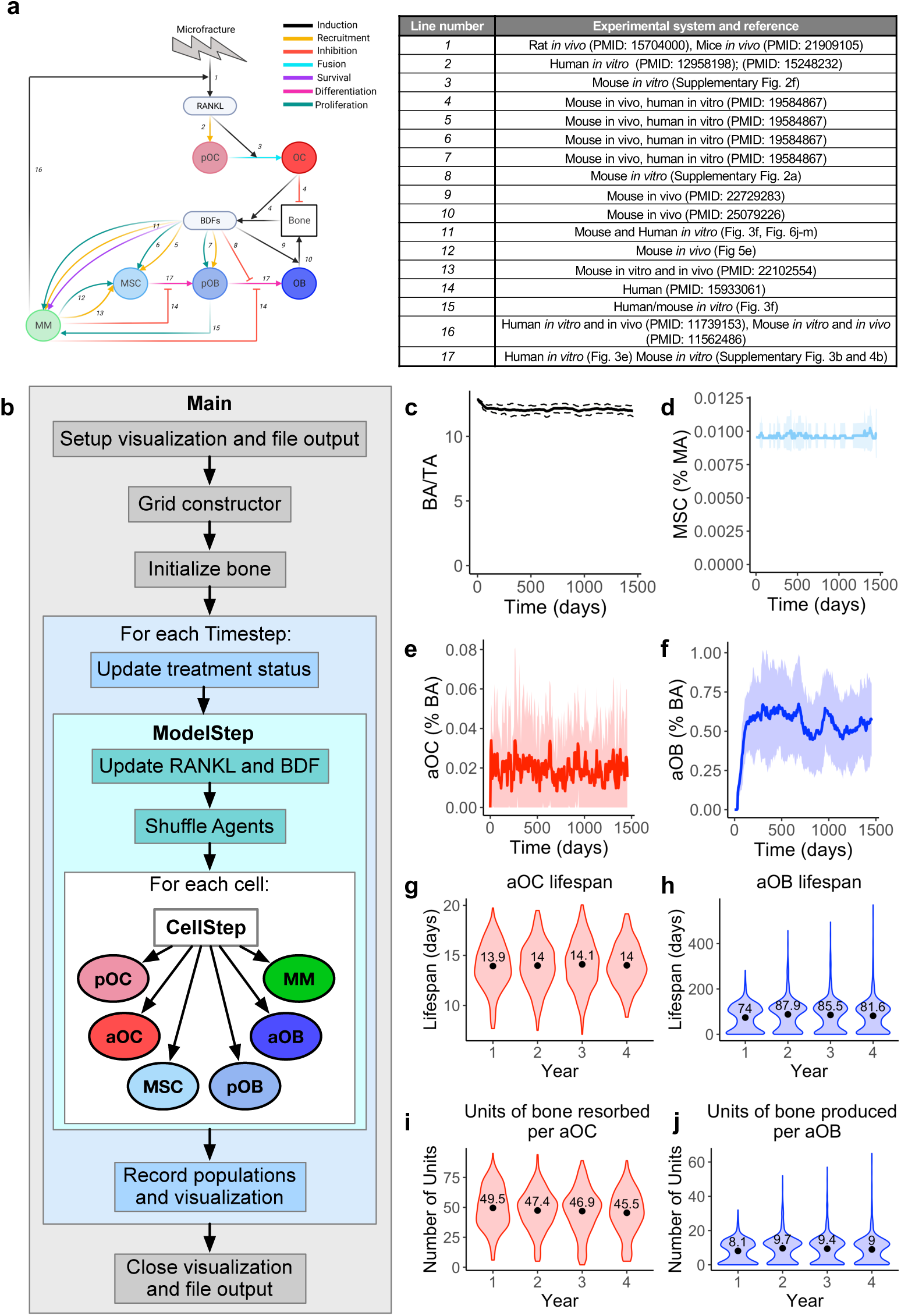
HCA captures key features of normal bone remodeling. **a.** Interaction diagram (left) between cell types in the HCA and factors such as BDFs and RANKL (created with biorender.com). Table of references and experimental systems in which results were obtained (right). **b.** HCA flow diagram adapted from Bravo et al. (2020)^6^. **c.** HCA model outputs of bone area to total area (BA/TA) **d.** HCA model outputs of MSC content. **e.** HCA model outputs of osteoclast numbers, and osteoblast numbers. **f-g**, Mean lifespan of osteoclasts **(f)** and osteoblasts **(g)** over 4 years of normal bone homeostasis. **h,** Mean units of bone resorbed per osteoclast each year of simulation. **i,** mean units of bone produced per osteoblast over course of the 4-year simulation.

**Supplementary Fig. 2.**
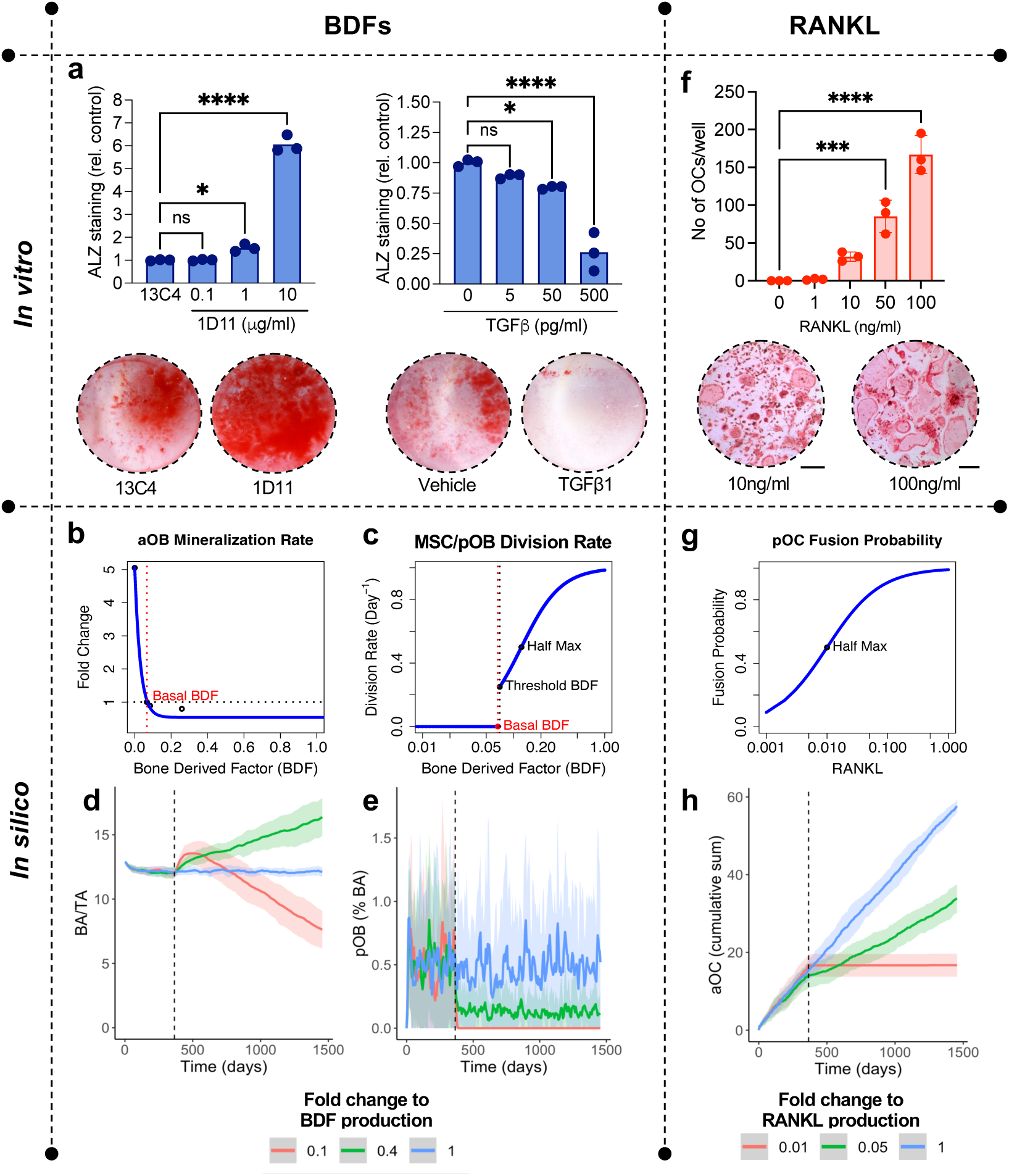
Inhibition of bone microenvironment cytokines in the HCA. **a,** The preosteoblast cell line MC3T3-E1 was cultured in osteogenic supplement with indicated concentrations of either isotype control antibody 13C4 or anti-TGF-β neutralizing antibody 1D11 (left) or TGF-β1 (right) for 14 days. Alizarin red was used to identify mineralized area. Results are normalized to isotype/vehicle-treated controls. Representative images of alizarin staining from control and highest concentrations (below). **b-c**, Plots of functional forms used to represent the fold change to osteoblast mineralization rate (**b**) and MSC/preosteoblast division rate (**c**). The black dots (**b**) represent the data from the *in vitro* experiments with the MC3T3-E1 cells that were used to fit the exponential decay function, as described in section 1.1 of supplementary methods. The red dot (**c**) represents the lack of MSC/preosteoblast division rate when BDF is at a basal level (dotted red line), and the black dot (**c**) represents the division rate when BDF is at the threshold level for proliferation (dotted black line). Another black dot highlights the half maximum division rate defined by the Hill function. * p<0.05 ** p<0.01, *** p<0.001, **** p<0.0001.

**Supplementary Fig. 3.**
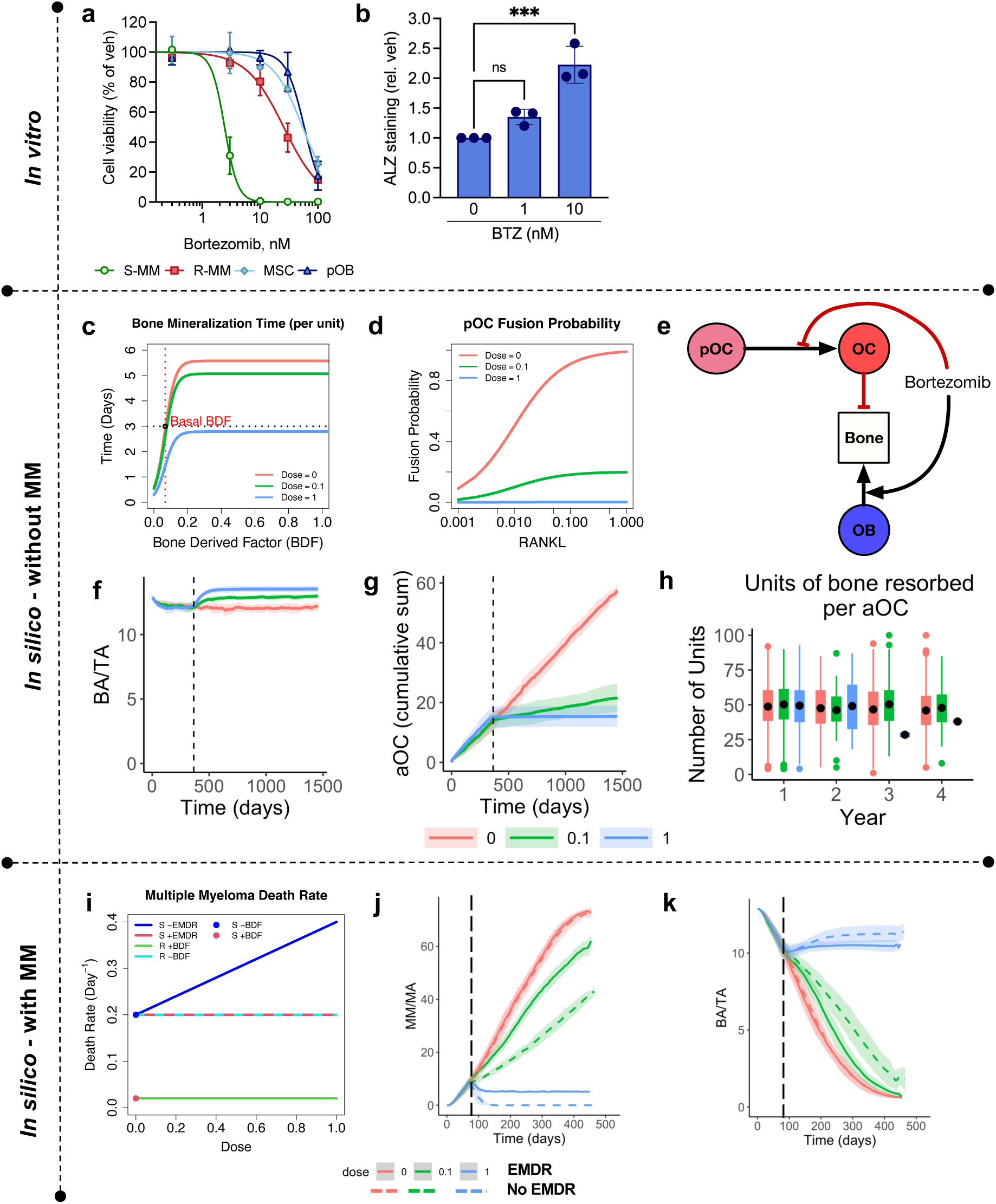
The impact of bortezomib (BTZ) on cells of the myeloma bone microenvironment *in vitro* and *in silico*. **a**, Sensitive U266 (S-MM) and resistant PSR (R-MM) myeloma, murine mesenchymal stromal cells (MSCs) and the preosteoblast cell line MC3T3-E1 (preosteoblast) were treated with varying concentrations of BTZ. Viability was measured after 72 hours by MTT assay. Results are normalized as percentage of control. **b**, the preosteoblast cell line MC3T3-E1 were cultured in osteogenic supplement with indicated concentrations of BTZ. Alizarin red was used to identify mineralized area. Results are normalized to vehicle-treated controls. **c-d**, Plots of functional forms used to represent the osteoblast mineralization time (**c**) and the probability of preosteoclast fusion (**d**) with indicated concentrations of bortezomib. **e**, Interaction diagram of normal bone cells with bortezomib. Red line denotes inhibition, black like indicates stimulation. **f-h**, Computational outputs of bone area to total area ratio (**f**), cumulative number of osteoclasts (**g**), and units of bone resorbed per osteoclast (**h**) following treatment of normal bone model with indicated concentrations of bortezomib. Treatment begins on day 365 (black dotted line) and is applied continuously for 3 years. **i**, Plot showing S-MM and R-MM death rate as a function of bortezomib with/without (+/−) BDF and +/− EMDR. **j**, Model outputs of MM growth with high and low doses of bortezomib with/without EMDR. Treatment begins when MM/MA=10% (black dotted line) and is applied continuously until MM/MA=20%. **k**, Flow chart outlining decisions MM cells make under BTZ treatment *** p<0.001.

**Supplementary Fig. 4.**
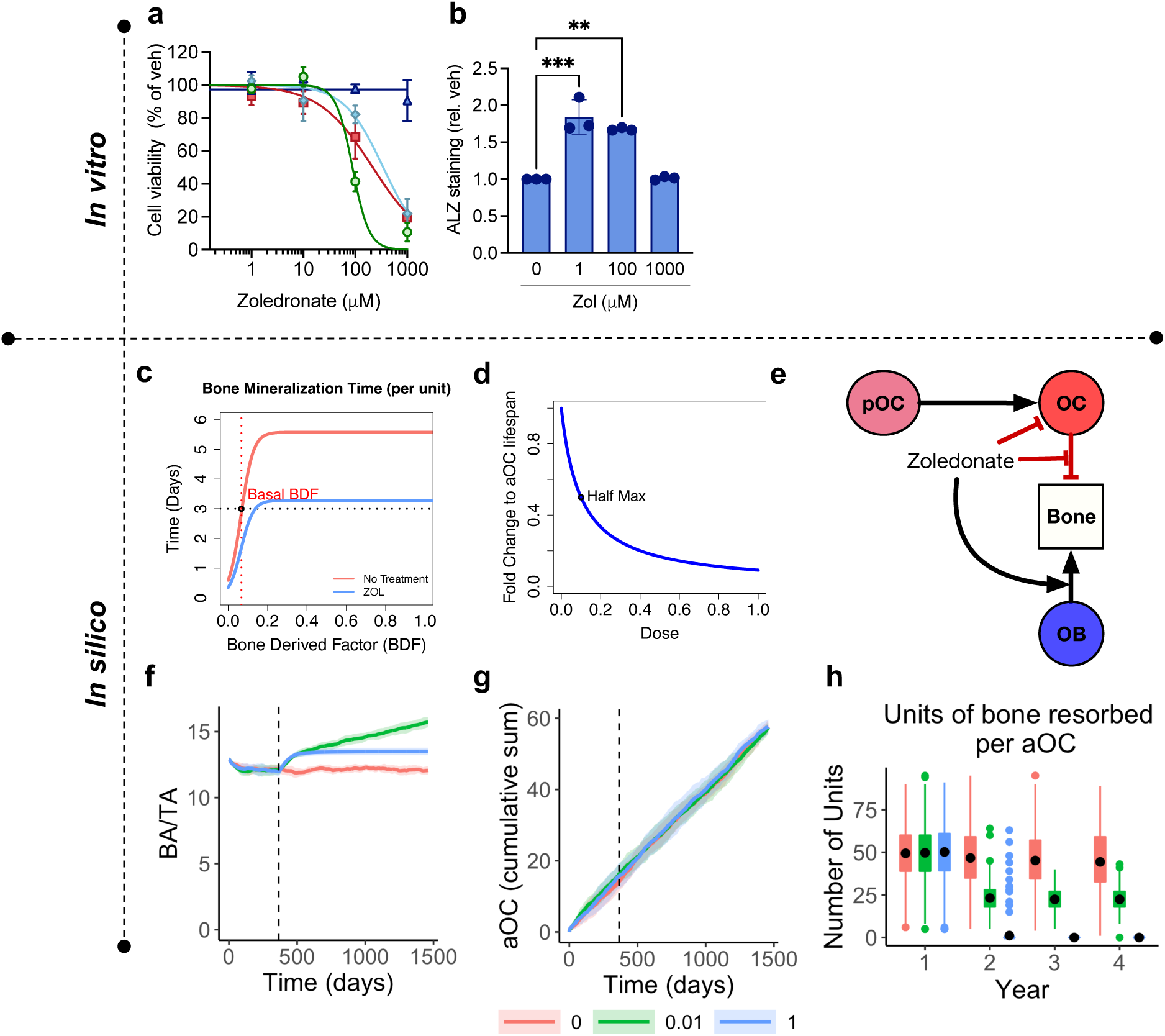
The impact of zoledronate (ZOL) on cells of the myeloma bone microenvironment *in vitro* and in *silico*. **a**, Sensitive U266 (S-MM) and resistant PSR (R-MM) myeloma, murine mesenchymal stromal cells (MSCs) and the preosteoblast cell line MC3T3-E1 (preosteoblast) were treated with varying concentrations of zoledronate (ZOL). Viability was measured after 72 hours by MTT assay. Results are normalized as percentage of control. **b**, The preosteoblast cell line MC3T3-E1 were cultured in osteogenic supplement with indicated concentrations of ZOL. Alizarin red was used to identify mineralized area. Results are normalized to vehicle-treated controls. **c-d**, Plots of functional forms used to represent the osteoblast mineralization time (**c**) and the fold change to osteoclast lifespan (**d**) with indicated concentrations of ZOL. **e**, Interaction diagram of normal bone cells with ZOL. Red line denotes inhibition, black like indicates stimulation. **f-h**, Computational outputs of bone area to total area ratio (**f**), cumulative number of osteoclasts (**g**), and units of bone resorbed per osteoclast (**h**) following treatment of normal bone model with indicated concentrations of ZOL. Treatment begins on day 365 (black dotted line) and is applied continuously for 3 years, *** p<0.001, **** p<0.0001.

**Supplementary Fig. 5.**
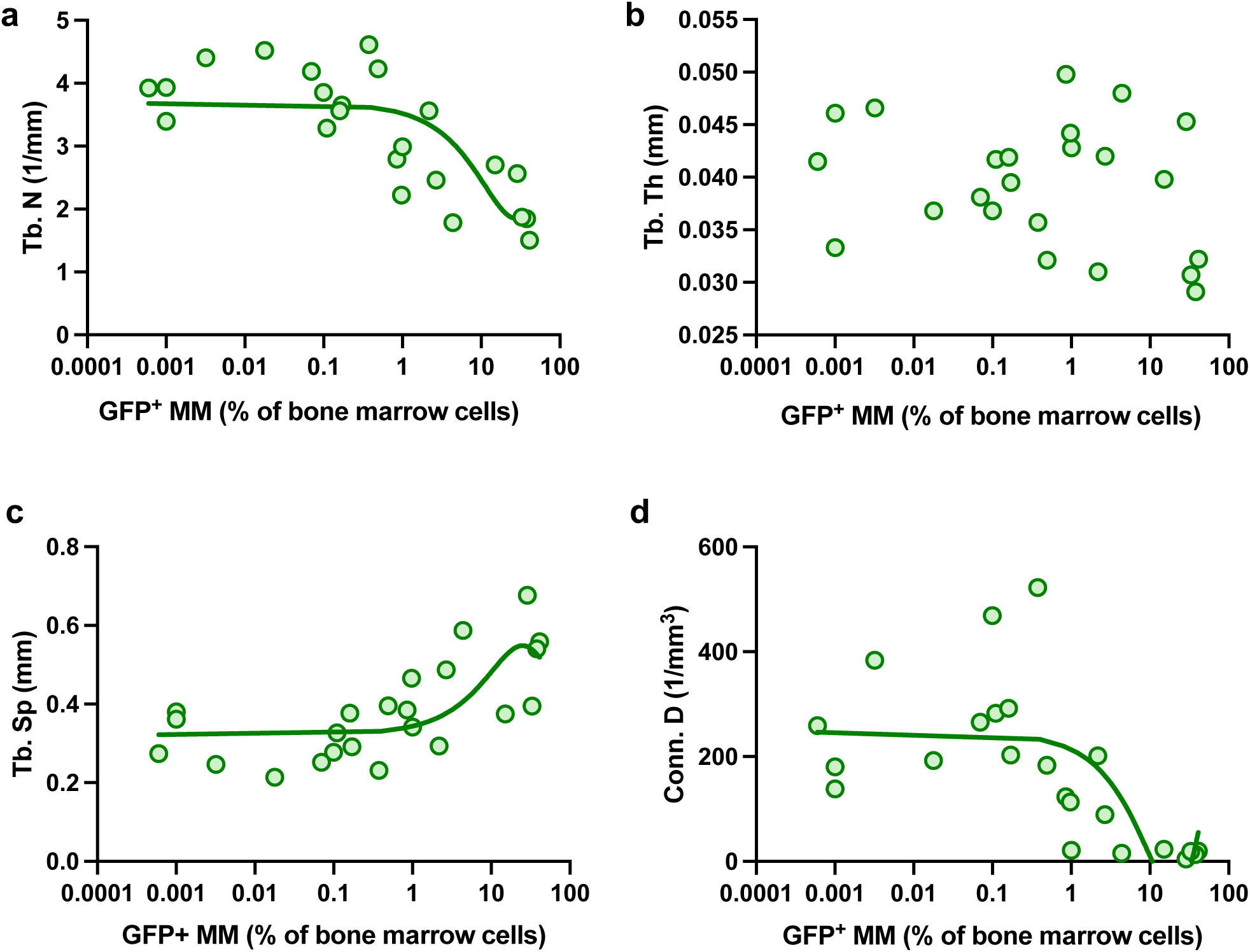
Myeloma alters bone microarchitecture *in vivo*. **a-d**, Tumor burden in tibia and femur was assessed by GFP positivity by flow cytometry in U266-bearing mice. High resolution microCT was used to assess trabecular number (Tb.N; **a**), thickness (Tb.Th; **b**) and spacing (Tb.Sp; **c**) and degree of connectivity (Conn.D; **d**).

**Supplementary Fig. 6.**
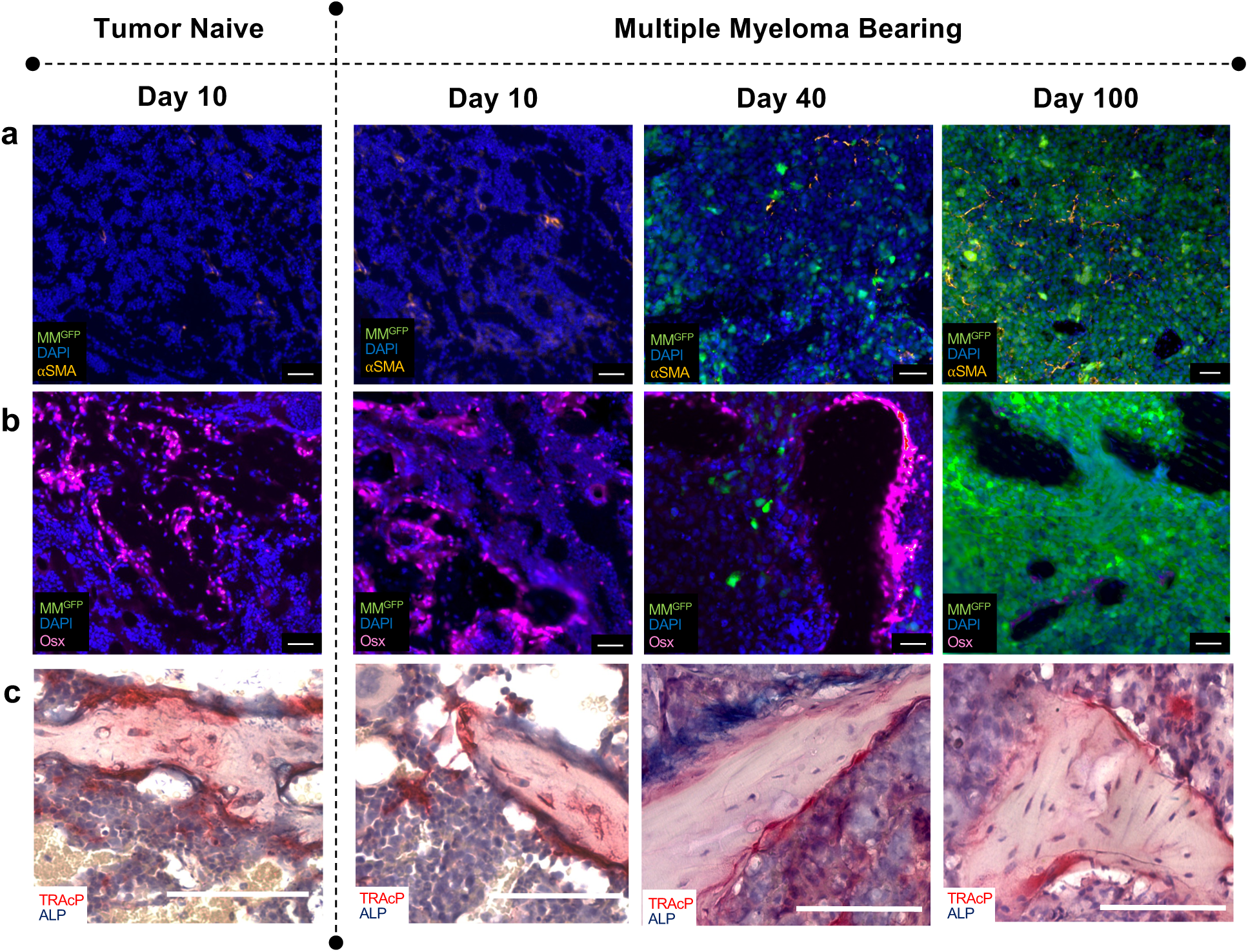
Multiple myeloma alters bone marrow microenvironment. **a-b**, Tumor naïve or U266GFP-bearing mice were sacrificed on indicated days to identify and localize *α*SMA+ mesenchymal stromal cells (gold; **a**), osterix+ preosteoblasts (pink; **b**) and GFP+ U266 cells (green; **a-b**) by immunofluorescence. Scale bars = 50µm **c**, Brightfield images of TRAcP+ multinucleated osteoclasts (red) and ALP+ cuboidal bone-lining osteoblasts (blue). Scale bars = 100µm. Images correspond with quantifications from Figure 6.

**Supplementary Fig. 7.**
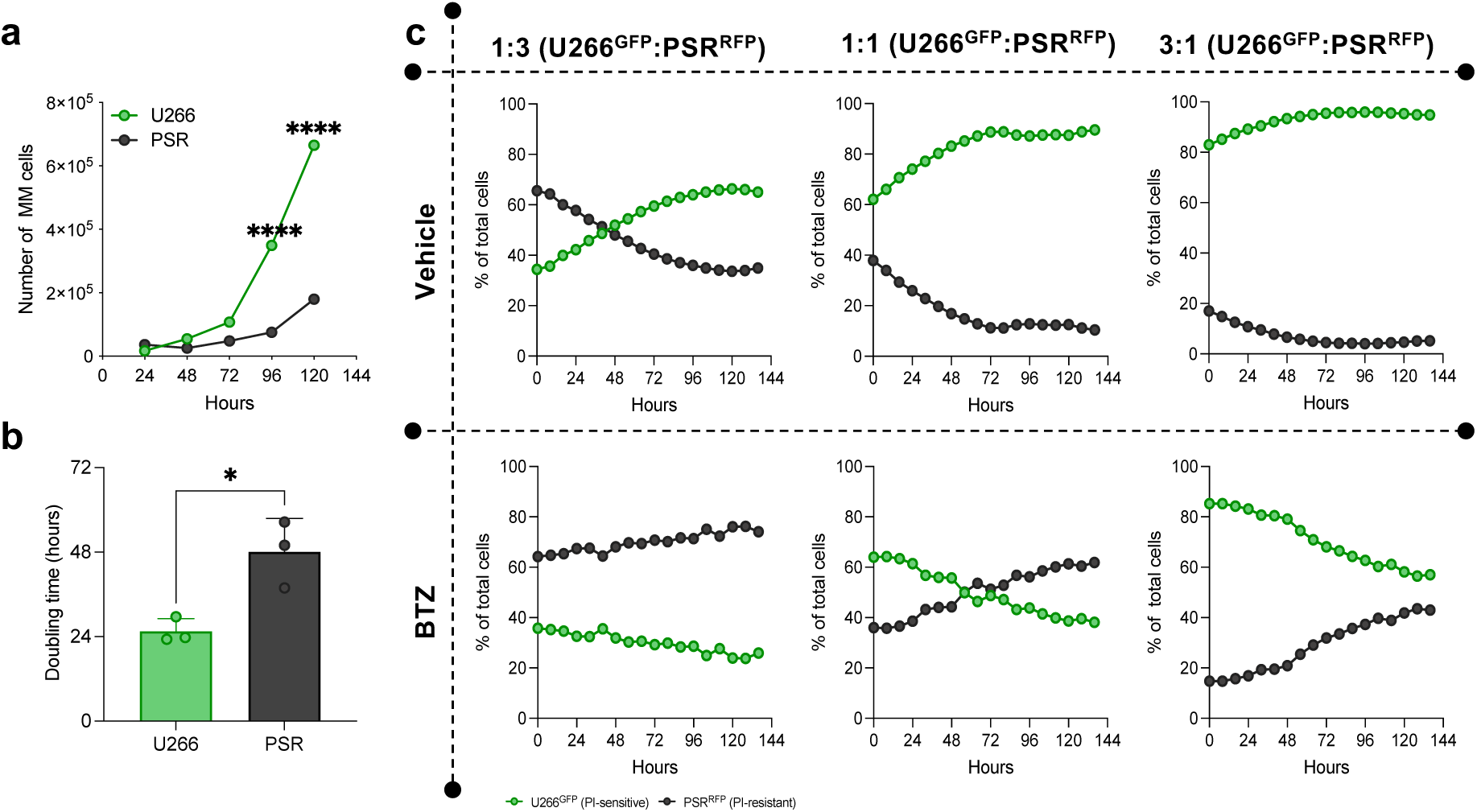
PI-sensitive MM cells outgrow their PI-resistant counterparts under normal growth conditions. **a-b**, PI-sensitive U266 and PI-resistant PSR MM cell lines were grown in standard culture conditions, the number of cell number was recorded using trypan blue staining over 120h (**a**) and used to calculate doubling times (**b**). Proteasome inhibitor sensitive GFP expressing U266 MM cells were incubated at indicated ratios with their proteasome inhibitor RFP expressing counterparts (PSR). The percentage of each population under vehicle control conditions was monitored over time using real time imaging microscopy (Incucyte). In parallel the impact of BTZ (4nM) on the outgrowth of the populations was observed revealing the emergence of the resistant population over time.

**Supplementary Fig. 8.**
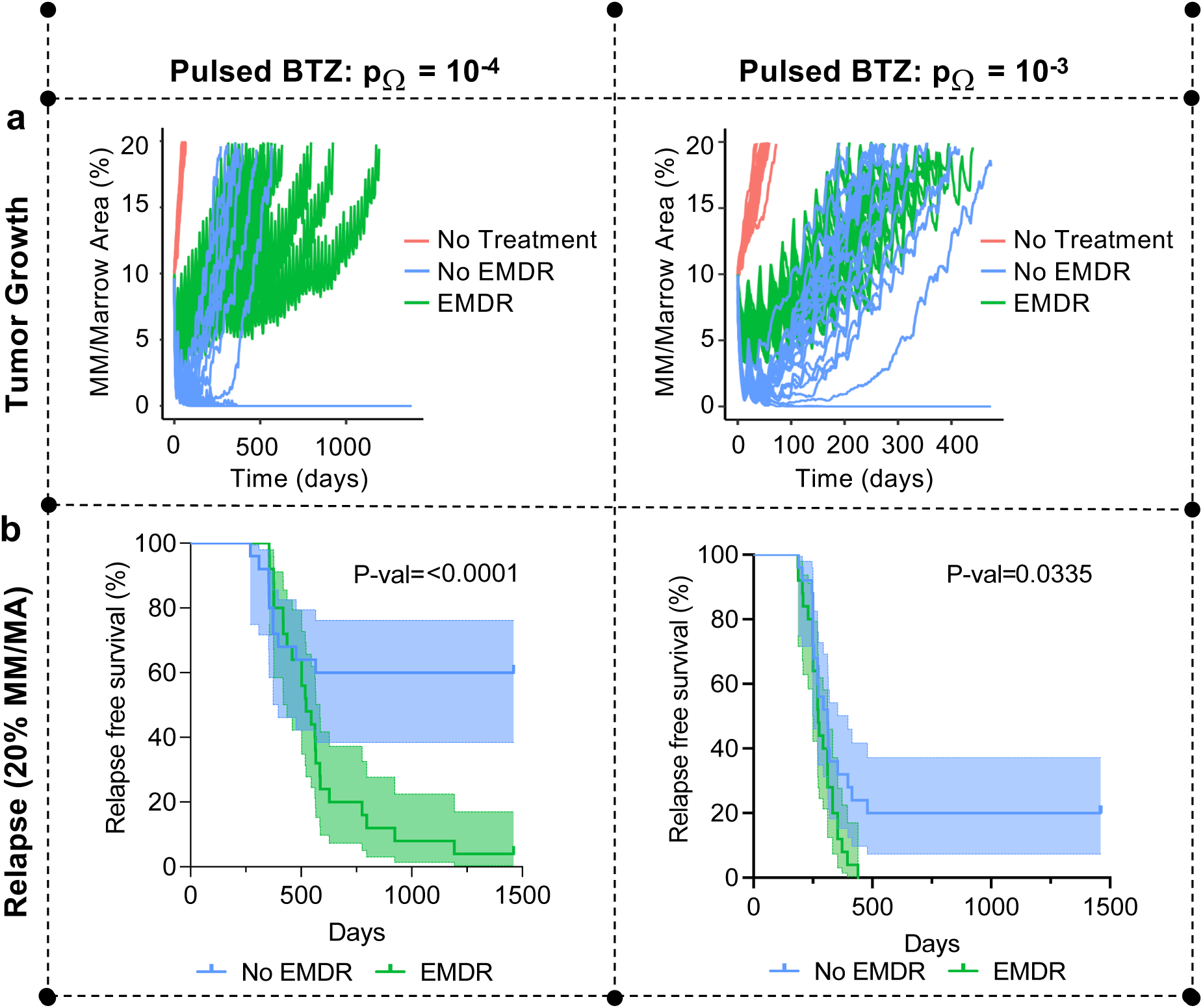
EMDR increases rate of myeloma relapse and tumor heterogeneity under pulsed bortezomib treatment. Pulsed BTZ treatment (2 weeks on therapy, 1 week off) was applied to HCA model when MM burden reached 10% of the marrow. **a-b**, Tumor growth (**a**) for individual simulations (n = 25 per condition) and Kaplan Meier plots of rate of relapse (**b**) for pulsed BTZ therapy were calculated when p_Ω_ = 10^−4^ (left column) and p_Ω_ = 10^−3^ (right column).

**Supplementary Fig. 9.**
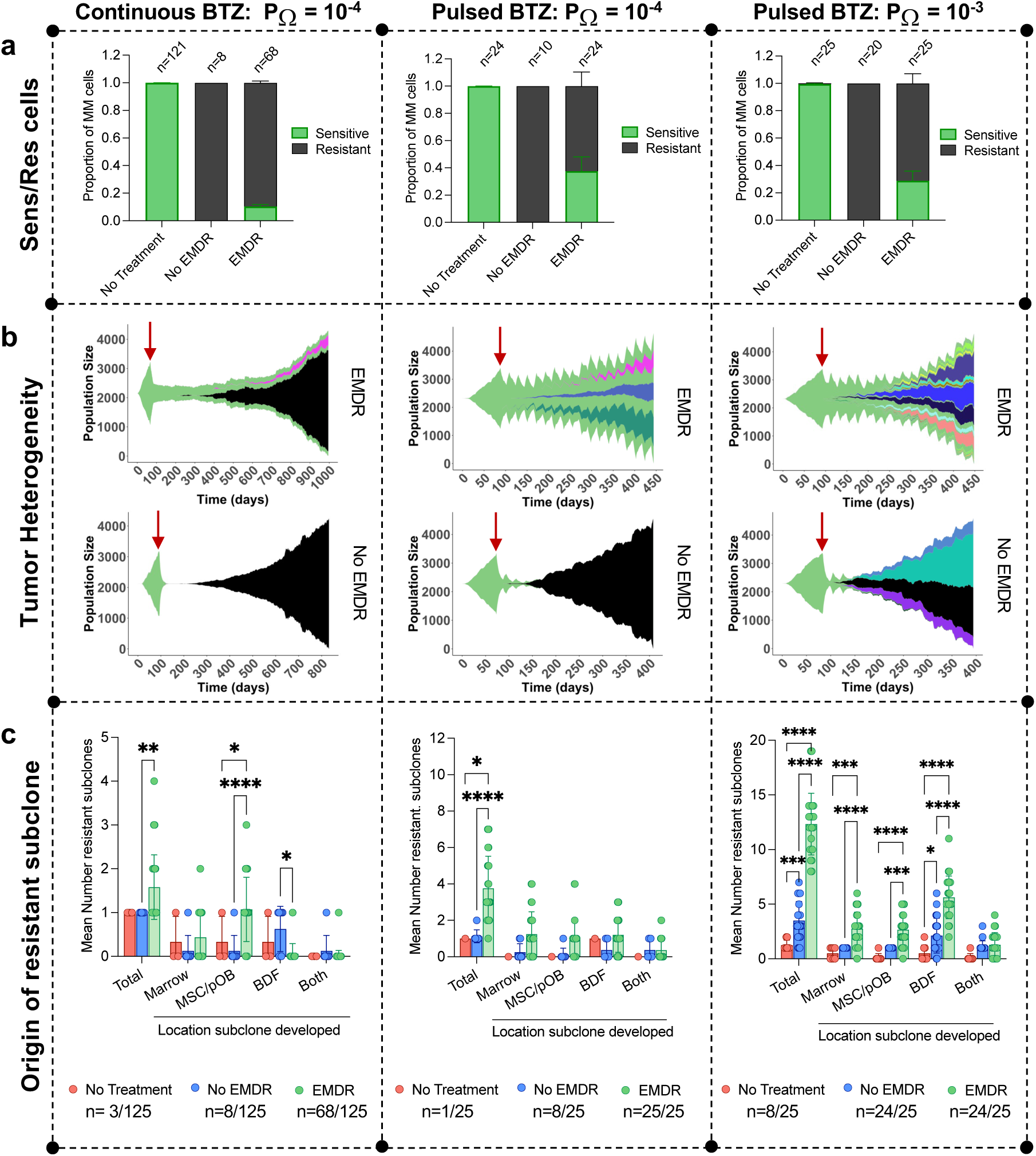
EMDR increases tumor heterogeneity under continuous and pulsed bortezomib treatment. Continuous BTZ treatment (left column) or Pulsed BTZ treatment (2 weeks on therapy, 1 week off; middle and right column) were applied to HCA model when MM burden reached 10% of the marrow. **a-c**, The proportion of sensitive/resistant MM cells at end point (MM = 20% of the marrow; **a**) and tumor heterogeneity (**b-c**) were assessed with two resistance probabilities (left and middle columns, p_Ω_ = 10^−4^, right column p_Ω_ = 10^−3^) in the presence or absence of EMDR. **b**, Muller plots indicating resistant subclones arising in tumor, created using EvoFreq^62^. Each color represents a unique subclone. Red arrow indicates start of treatment. **c**, Mean number of resistant sub-clones arising in tumors that reached 20% and contained resistant subclones (MM/MA) following continuous or pulsed BTZ treatment at different locations within the BME with/without EMDR. ‘Both’ refers to resistant subclones that arise close to MSCs and BDFs. Overall, EMDR leads to significantly higher numbers of resistant subclones.

**Supplementary Fig. 10.**
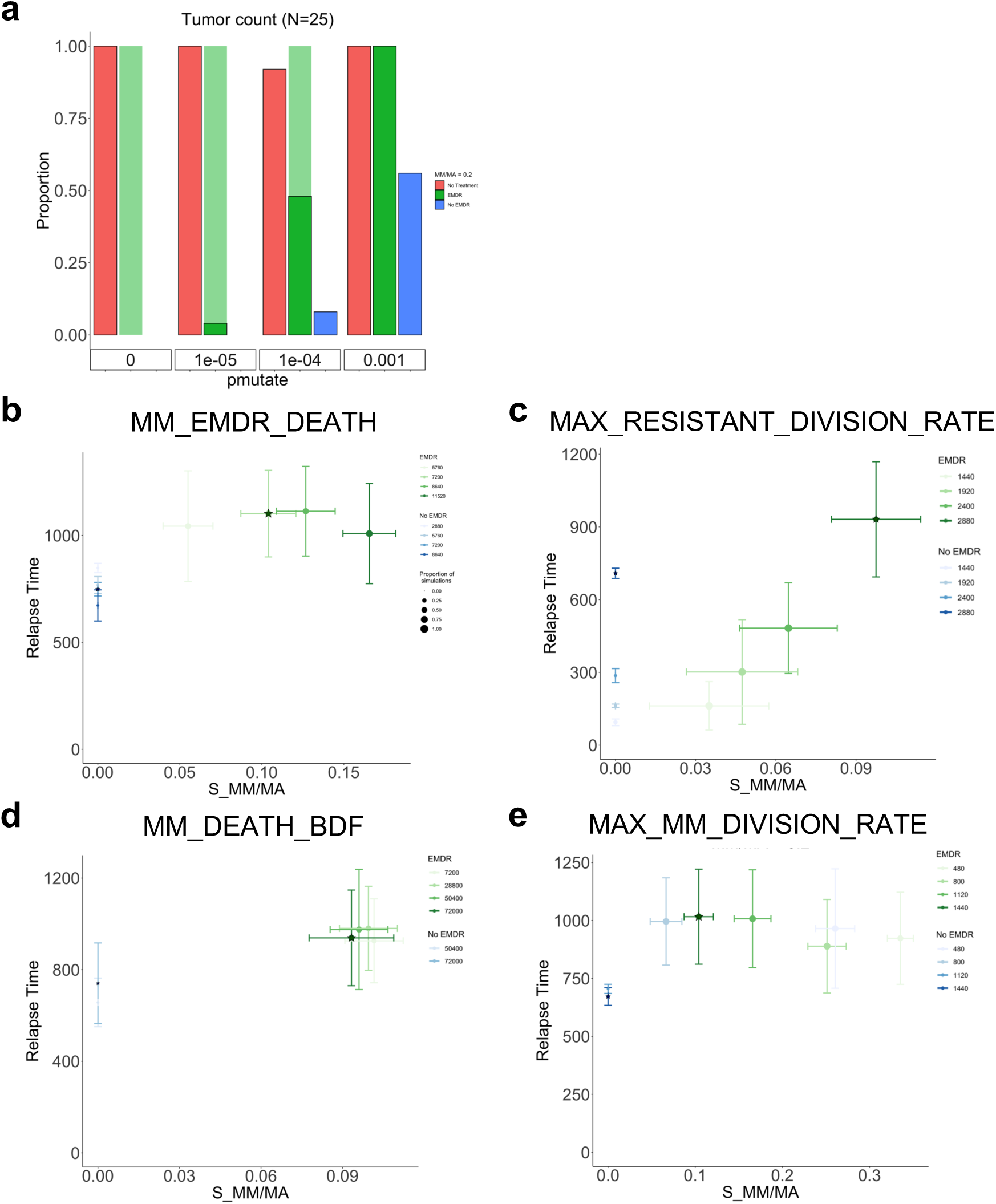
The impact of select parameter values on key model outputs. Continuous BTZ treatment was applied to HCA model when MM burden reached 10% of the marrow. **a.** Proportion of tumors that survived (lighter shade) or relapsed (darker shade) was assessed as resistance probability increased. **b-e**. Select parameters were varied to assess how the relapse time (when MM burden reached 20% of the marrow) and proportion of sensitive cells at endpoint changed due to parameter values (p_Ω_ = 10^−4^). MM_EMDR_DEATH is the lifespan of sensitive cells (+ BTZ) protected by EMDR (**b**). MAX_RESISTANT_DIVISION_RATE is the maximum division time of resistant cells (+/− BTZ) (**c**). MM_DEATH_BDF is the lifespan of sensitive cells (-BTZ) or resistant cells (+/− BTZ) in the presence of BDF (**d**). MAX_MM_DIVISION_RATE is the maximum division time of sensitive cells (+/− BTZ) in the presence of MSC/preosteoblast (**e**). The star marker indicates the outputs corresponding to the baseline parameter value. The circular marker indicates the mean value of the outputs with error bars representing the standard deviation of the simulations (n = 25); the size of the circular marker increases as the proportion of simulations that reached the endpoint increased.

**Supplementary Video 1. HCA model recapitulates normal bone remodeling.**

HCA simulation of normal bone remodeling, showing agent grid (top) and PDE grids with scaled concentration of RANKL (bottom left) and bone derived factors (BDF, bottom right) over one year. Each bone remodeling event is initiated through a small microfracture, which creates a signal to osteoblast lineage cells to express RANKL. Preosteoclasts migrate to the bone remodeling site and undergo fusion in response to RANKL to become a fully formed osteoclast. As the osteoclast resorbs the bone, BDF is released from the bone matrix which promotes the proliferation of preosteoblasts. During the reversal period, BDF continues to be released at a lower level, causing the preosteoblasts to migrate and attach to the eroded surface. Low BDF promotes the differentiation of preosteoblasts into bone-forming osteoblasts, which continue to build bone until the amount of bone that was resorbed is replaced. The bone formation phase is then followed by a period of quiescence until bone remodeling is re-initiated at the given location.

YouTube link: https://www.youtube.com/watch?v=UFpXzLNFMzo

**Supplementary Video 2. HCA model captures key steps of myeloma-bone vicious cycle.**

HCA simulation of the myeloma-bone vicious cycle, showing agent grid (top) and PDE grids with scaled concentration of RANKL (bottom left) and bone derived factors (BDF, bottom right) over one year. Each simulation is initialized with a single MM cell after the formation of the first osteoclast. MM increases the number of osteoclasts which drives bone resorption and the release of BDFs. BDFs and MSCs that are recruited in response to MM contribute to the growth and survival of MM cells (light green cells), causing MM cells to further expand in the bone marrow. At the same time, MM cells inhibit the differentiation of MSCs and preosteoblasts, preventing the formation of bone and further driving osteolytic bone disease.

YouTube link: https://www.youtube.com/watch?v=NFdpWp2l1wg

**Supplementary Video 3. EMDR increases minimal residual disease.**

HCA simulation of continuous treatment with bortezomib (BTZ), showing agent grid (top) and PDE grids with scaled concentration of RANKL (bottom left) and bone derived factors (BDF, bottom right). Treatment begins when multiple myeloma (MM) cells have reached 10% of the bone marrow (the starting point of the supplemental video) and continues until the MM burden doubles. After treatment begins, MM cells have a probability of resistance (p_Ω_) during cell division that causes resistance to BTZ. When p_Ω_ =10^−4^, EMDR (middle) protected a portion of sensitive cells, enabling them to acquire resistance and drive tumor relapse. When EMDR is absent (right), the tumor went extinct.

YouTube link: https://www.youtube.com/watch?v=fnBuxXvWWXw

**Supplementary Video 4. Higher resistance probability increases tumor relapse rates in absence of EMDR.**

HCA simulation of continuous treatment with bortezomib (BTZ), showing agent grid (top) and PDE grids with scaled concentration of RANKL (bottom left) and bone derived factors (BDF, bottom right). Treatment begins when multiple myeloma (MM) cells have reached 10% of the bone marrow (the starting point of the supplemental video) and continues until the MM burden doubles. After treatment begins, MM cells have a probability of resistance (p_Ω_) during cell division that causes resistance to BTZ. When p_Ω_ =10^−3^, EMDR (middle) protected a portion of sensitive cells, enabling them to acquire resistance and drive tumor relapse. When EMDR is absent (right), the high resistance probability prevents tumors from going extinct before the acquisition of intrinsic resistance and also drives tumor relapse.

YouTube link: https://www.youtube.com/watch?v=EJUOy0yepWw

